# Psychedelics Relax Priors and Reshape Orbitofrontal Dynamics

**DOI:** 10.1101/2025.09.18.677110

**Authors:** C. Delgado-Sallent, S. A. Ahmed, A. Khawaja-Lopez, B. S. Lee, R.A. Senne, B. B. Scott, S. Ramirez

## Abstract

Psychedelics such as psilocybin and ketamine are gaining attention as rapid-acting treatments for psychiatric disorders, yet the mechanisms by which they alter cognition remain unclear. A key hypothesis—the REBUS model—proposes that psychedelics relax high-level priors, allowing bottom-up sensory information to exert greater influence over perception and behavior. Here, we test this model in mice performing a free-response perceptual decision-making task that disambiguates prior-driven and sensory-driven decision strategies. Acute administration of psilocybin or ketamine significantly slowed decision times and improved accuracy. Behavioral modeling that combined drift diffusion and GLM-HMM frameworks revealed that these changes were mediated by increased decision thresholds and a marked shift into sensory-engaged cognitive states. Whole-brain c-Fos mapping identified a distributed decision-making network, with psychedelics selectively modulating cortical and subcortical nodes. Calcium imaging in the orbitofrontal cortex (OFC)—a key region for integrating priors and sensory inputs—revealed preserved decision-related selectivity under psychedelics, while exhibiting reduced neuronal correlations—population-level signatures of weakened top-down influence and relaxed priors. Together, these results provide circuit-level support for the REBUS model, showing that psychedelics reconfigure brain-wide and local dynamics to promote more deliberate, flexible, and sensory-driven decision policies.

## Introduction

Psychedelics like psilocybin and ketamine have shown great potential in treating psychiatric disorders, often producing long-lasting effects after a single dose^1,2^. Psychedelic compounds produce profound alterations in perception and cognition, hypothesized to stem from a relaxation of the brain’s normally constraining prior beliefs. The “Relaxed Beliefs Under Psychedelics” (REBUS) model proposes that psychedelics reduce the influence of high-level priors in the brain’s hierarchical predictive processing architecture. In effect, top-down influences are diminished, allowing bottom-up sensory information to drive perception more freely^3,4^. This mechanistic framework has been invoked to explain the hallucinatory experiences and cognitive flexibility reported in psychedelic states, but it remains untested in causal, circuit-level experiments.

Perceptual decision-making offers a principled framework for probing how psychedelics alter the balance between prior expectations and sensory evidence^5–7^. Recent advances in task design use carefully controlled, noisy stimuli to quantify how behavior emerges from sensory input, internal decision policies, and prior beliefs^8^. In particular, tasks that allow subjects to freely balance speed and accuracy expose how internal policies govern the trade-off between rapid, bias-driven choices and slower, evidence-based inference—key predictions of the REBUS model^3^. These paradigms are readily implemented across species, enabling fine-grained measurement of neural dynamics in rodent models^9,10^.

To test this, we administered psilocybin (a classic 5-HT2A agonist psychedelic) or ketamine (an NMDA receptor antagonist with psychedelic-like effects) to mice performing a perceptual integration task designed to probe the balance between prior expectations and sensory evidence^9,10^. We combined behavioral analysis with neural measurements across multiple scales to assess whether these drugs shift the balance between prior-driven and sensory-driven decision processes. Behaviorally, we found that psychedelics weakened prior-driven biases and enhanced reliance on sensory evidence, aligning with REBUS model predictions. Whole-brain c-Fos mapping identified a conserved network for decision-making, with notable modulation in the orbitofrontal cortex (OFC)—a region implicated in value-guided behavior, belief updating, and encoding task structure^11,12^. Cellular-resolution calcium imaging revealed that the OFC preserves decision-related selectivity under psychedelics, while exhibiting reduced neuronal correlations—population-level signatures of weakened top-down influence and relaxed priors.

By examining behavior, circuit-wide activation patterns, and OFC neuronal ensemble activity, we provide evidence for core predictions of the REBUS model – namely that psychedelics relax prior beliefs, bolster bottom-up sensory influence, and increase the brain’s cognitive flexibility – in an animal model of perceptual decision-making.

## Results

### Sensory Evidence and Priors Shape Decision Strategies during a Free-Response Perceptual Integration Task

To investigate how sensory evidence and priors shape decisions, we trained mice on a free-response perceptual integration task^9,10^. Mice initiate each trial with a nose poke, then observe a rapid series of randomly timed light flashes from two locations (left vs. right). On each trial, one side is designated “correct” (drawn randomly), with a higher flash probability (e.g., 80% vs. 20%, see Methods). In this task, mice can freely decide when they have enough evidence to nosepoke either the left or right port, at which point stimulus presentation ceases (Figure 1A, See methods). This design allowed mice to dynamically balance the trade-off between additional evidence accumulation and decision commitment, providing insight into their reliance on sensory information versus prior expectations (Figure 1A-B). In this framework, we treat decision biases—like side preference or win-stay behavior—as proxies for task-specific priors, reflecting structured expectations.

**Figure 1.**
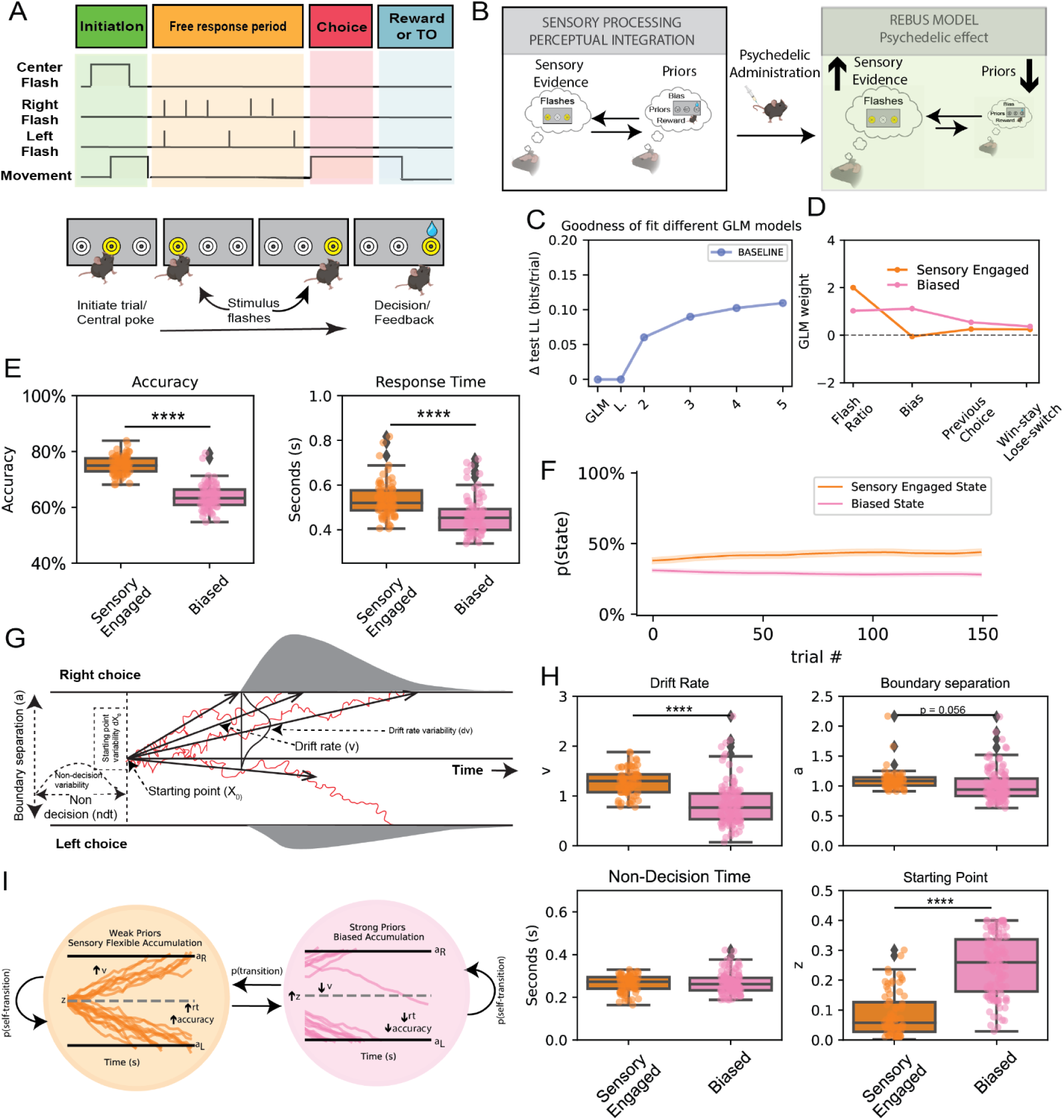
A free-response evidence accumulation task reveals distinct decision-making states modulated by sensory evidence and priors. (A) Schematic of the behavioral task. Mice initiate trials by poking a central port, then receive randomly timed left/right visual flashes until they make a decision by choosing one of the side ports. The task is free-response, allowing animals to freely decide when to respond. (B) Conceptual framework linking the task to the REBUS model. Mice can rely on either bottom-up sensory evidence or top-down priors (e.g., biases or reward history). Psychedelic administration is proposed to reduce the influence of priors, increasing sensitivity to sensory evidence. **(C)** Model comparison showing that a 3-state Generalized Linear Model–Hidden Markov Model (GLM-HMM) best captures behavioral data, with a sharp improvement in model fit over simpler models. **(D)** GLM weights for behavioral regressors in the sensory engaged and biased states. Sensory evidence (flash ratio) carries more weight in the sensory engaged state, whereas priors (e.g., previous choice and win-stay/lose-switch strategies) dominate the biased state. **(E)** Behavioral performance stratified by state. Mice in the sensory engaged state show significantly higher accuracy and slower response times compared to the biased state, indicating a more deliberate accumulation of evidence. **(F)** State occupancy over trials shows relatively stable engagement of each strategy across time. **(G)** Drift Diffusion Model (DDM) schematic illustrating key latent variables: drift rate (v), boundary separation (a), starting point (z), and non-decision time (ndt). **(H)** DDM parameter comparisons between states. The sensory engaged state is associated with significantly higher drift rates and a lower starting point bias, and trends toward higher boundary separation (p = 0.056), reflecting greater reliance on sensory evidence and less bias. **(I)** Conceptual model of decision state transitions. In the sensory engaged state, decisions are guided by flexible, sensory-driven accumulation. In the biased state, strong priors dominate accumulation, leading to faster but less accurate decisions. Notations: * = p<0.05, ** = p<0.01, *** = p<0.005, **** = p<0.001.

We reasoned that in some trials, decisions might rely more on *bottom-up* sensory evidence (“sensory-engaged” state), whereas in others, *top-down* biases like choosing the preferred side or the side of last reward might dominate (“biased” state). While not all biases represent hierarchical priors in the REBUS sense, they operationally capture pre-evidence influences on choice. The REBUS model predicts psychedelics shift behavior toward the sensory-engaged state by reducing prior influence (Figure 1B). To formalize these latent states, we fit a Hidden Markov Model variant of a Generalized Linear Model (GLM-HMM)^13^. We compared models with different state numbers, finding that a three-state model best explained choice behavior. These states corresponded to: (1) a *sensory-engaged* state, where choices depended strongly on the current flash evidence; (2) a *left-biased* state; and (3) a *right-biased* state. For visualization purposes, left/right-biased states were grouped together, as both reflect heavy reliance on priors (Figure 1C; Supplementary Figure 1). In the biased state, choices were influenced by prior biases (e.g., win-stay or side bias) rather than incoming evidence, whereas in the engaged state, current sensory evidence dominated. This separation was validated by state-dependent performance differences: sensory-engaged bouts yielded higher accuracy than biased bouts, confirming that real-time evidence used improved performance (*T = 255.65, p < 0.0005*; Figure 1D-E). Furthermore, response times were significantly slower in the sensory engaged state, suggesting that prioritizing sensory information over priors requires additional processing time to accumulate sufficient evidence before making a decision *(T = 46.83, p < 0.0005*; Figure 1E).

To assess the stability of decision-making strategies, we quantified how long mice remained in each GLM-HMM-defined state. Transition analyses revealed that once animals entered a given state—either sensory-engaged or biased—they tended to persist across trials (Figure 1F, Supplementary Figure 1A). This persistence suggests that decisions are governed by consistent internal strategies, aligning with REBUS model predictions that weakening top-down priors should promote stable engagement with bottom-up sensory evidence, rather than impulsive switching between strategies^13^.

To further characterize the decision-making process, we applied Drift Diffusion Modeling (DDM) to estimate latent cognitive parameters, an approach that aims to characterize shifts in reliance on prior expectations versus sensory evidence across distinct behavioral states (Figure 1G)^5,14^. Drift rate, which quantifies the strength of sensory evidence was significantly higher in the sensory engaged state (*T = 9.48, p < 0.0005*). Boundary separation was also slightly larger in engaged vs. biased states, meaning that in engaged states mice set a higher threshold, consistent with their longer response times (*T = 1.93, p = 0.056*). The starting point of the decision process was significantly shifted in the biased state, indicating a strong prior-driven influence on decision-making *(T = -20.47, p < 0.0005*). Non-decision time, which accounts for sensory encoding and motor execution, did not differ significantly between states, suggesting that observed differences were primarily due to cognitive rather than motor processes (Figure 1H).

Together, these findings demonstrate that this free-response perceptual integration task effectively captures and parametrizes the transitions between a sensory engaged state and a biased state (Figure 1I). This framework provides a valuable tool for investigating how psychedelics influence these decision-making states and whether they modulate transitions in accordance with the predictions of the REBUS model.

### Psychedelics Weaken Prior Influences and Enhance Perceptual Integration by Increasing Decision Thresholds

To investigate how psychedelics influence decision-making strategies, we administered varying doses of psilocybin (0.5, 1, and 3 mg/kg), ketamine (10, 30, 50, and 70 mg/kg), or saline to mice performing the free-response perceptual integration task.

We first analyzed accuracy and response times under psilocybin, ketamine, and saline conditions (Figure 2A). Psilocybin induced a dose-dependent increase in both accuracy and response times *(Accuracy: F = 13.59, p < 0.0005; RT: F = 24.77, p < 0.0005*). Ketamine produced a similar dose-dependent effect, where moderate doses (30–50 mg/kg) improved accuracy and response times, while lower (10 mg/kg) and higher doses (70 mg/kg) had no significant impact on performance (*Accuracy: F = 13.84, p < 0.0005; RT: F = 31.44, p < 0.0005*). Changes in performance were stable throughout the session with psilocybin, whereas ketamine’s effect peaked within the first 20 minutes before declining (Supplementary Figure 2A). In contrast, cocaine—used as a non-psychedelic control—did not affect performance, suggesting that the observed effects are specific to psychedelic compounds (Supplementary Figure 3).

**Figure 2.**
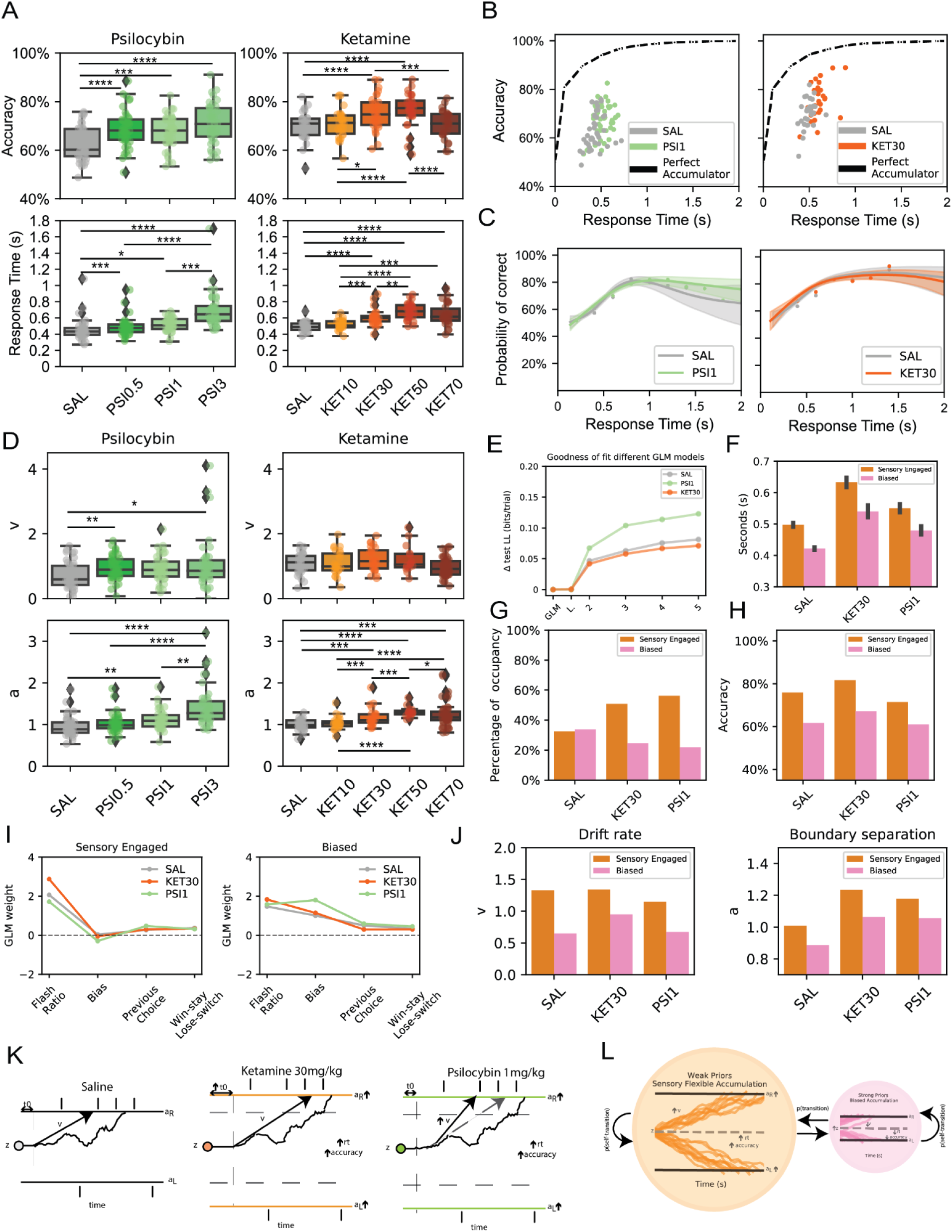
Psychedelics modulate perceptual decision-making by increasing decision thresholds and enhancing sensory engagement. **(A)** Behavioral performance across doses of psilocybin (left) and ketamine (right). Both compounds show dose-dependent increases in accuracy and response time at moderate doses, with performance impairments at higher doses for ketamine. **(B)** Speed-accuracy trade-off plots comparing psilocybin (left) and ketamine (right) to saline and a perfect accumulator model. Psychedelics shift the trade-off toward higher accuracy at longer decision times. **(C)** Probability of correct choice as a function of response time under saline and drug conditions. **(D)** Drift Diffusion Model (DDM) analysis of latent parameters. Drift rate (ν) and boundary separation (a) increase under psilocybin and ketamine increased boundary separation at medium doses, supporting enhanced evidence accumulation and more cautious decision thresholds. **(E)** Model comparison showing improved fit of GLM-HMM models with increasing complexity across conditions, especially under drug treatment. **(F)** Average response times in sensory engaged and biased states across saline, ketamine, and psilocybin groups. Psychedelics lengthen response time predominantly in the sensory engaged state. **(G)** Proportion of trials classified as sensory engaged or biased. Psychedelics increase the proportion of trials in the sensory engaged state. **(H)** Accuracy per decision state and treatment condition. Accuracy is higher in sensory engaged states across all conditions and is enhanced by psychedelics. **(I)** GLM weights for behavioral regressors in sensory engaged (left) and biased (right) states, showing stronger sensory evidence weighting in the engaged state and stronger prior-related influence in the biased state. Psychedelics reduce prior weighting and enhance sensory reliance. **(J)** DDM parameters split by state. Both psilocybin and ketamine increase drift rate and boundary separation in the sensory engaged state. **(K)** Schematic summary of neural trajectories under saline, ketamine, and psilocybin. Psychedelics increase decision thresholds (a) and improve accuracy by enhancing evidence accumulation. **(L)** Updated conceptual model: Psychedelics promote transitions to a sensory flexible accumulation state by flattening priors and increasing bottom-up processing, consistent with the REBUS model framework. Notations: * = p<0.05, ** = p<0.01, *** = p<0.005, **** = p<0.001.

To evaluate whether psychedelics improve decision-making by encouraging greater evidence accumulation, we examined the relationship between response time and accuracy—a key indicator of speed-accuracy trade-offs (Figure 2B). Under saline, mice exhibited a speed-accuracy trade-off pattern: faster responses tended to be less accurate. Under psilocybin (1 mg/kg) and ketamine (30 mg/kg), this relationship shifted upward. Mice waited longer on average, and those longer waits translated into even higher accuracy, suggesting that psychedelics encourage more evidence accumulation before decision commitment. Notably, despite slower decisions overall, the fundamental shape of the accuracy-vs-time curve remained intact, with peak accuracy for responses around ∼0.7–1.0 s for all conditions (Figure 2C).

Drift diffusion modeling reinforced these observations (Figure 2D, Supplementary Figure 2D–G). Under psychedelics, the dominant change was an increase in boundary separation (decision threshold). Both psilocybin and ketamine significantly raised the threshold (*Psilocybin: F = 20.20, p < 0.0005; Ketamine: F = 7.32, p < 0.0005*; Figure 2D), meaning mice required more accumulated evidence to respond. Psilocybin also modestly increased drift rate (ν), indicating faster evidence accumulation per unit time *(F = 3.57, p = 0.029*). Ketamine showed a similar trend at moderate doses, though to a lesser extent. Non-decision time was elevated only at high ketamine doses, suggesting possible motor slowing distinct from the cognitive effects *(F = 20.20, p < 0.0005*; Supplementary Figure 2E). Thus, psychedelics raised their internal threshold for terminating sampling.

To further characterize these effects, we used the three-state GLM-HMM framework introduced earlier (Figure 1C–H, Supplementary Figure 1), classifying trials into sensory engaged, left-biased, and right-biased states (with the biased states combined for visualization). Model comparisons showed that psychedelic administration and saline controls were fitted similarly by the model (Figure 2E).

Next, we examined how the improvement in accuracy under psychedelics was linked to decision-state transitions. Mice under psychedelics spent less time in biased states and more time in the sensory engaged state, which was associated with higher accuracy, longer response times, increased drift rate, and higher decision boundaries (Figure 2F–H, J). Notably, boundary separation was elevated in both sensory engaged and biased states under psychedelics, indicating higher decision thresholds independent of state (Figure 2J). Moreover, the sensory engaged state under ketamine produced the highest accuracy and the longest response times, demonstrating the strongest reliance on sensory evidence (Figure 2I). Finally, psilocybin and ketamine induced distinct temporal dynamics in state transitions throughout the session, which correlated with the temporal dynamics linked to the drug processing (Supplementary Figure 2A and H), suggesting that difference dose-response time dynamics in response to the different drugs.

Together, these findings support two key models of how psychedelics influence perceptual decision-making. First, psychedelics increase decision thresholds (Figure 2K), allowing for greater evidence accumulation before commitment, a mechanism that has been linked to reward-error encoding^15–17^. Second, psychedelics reduce prior expectations, shifting decision-making toward sensory-driven strategies (engaged-state; Figure 2L). These results align with the REBUS model, which proposes that psychedelics flatten hierarchical priors, thereby increasing reliance on real-time sensory information^3^. This framework provides a mechanistic explanation of how psychedelics influence cognition, with implications for their role in promoting flexible and adaptive sensory processing.

### Whole-brain c-Fos Mapping Identified a Consistent Neural Circuit for Perceptual Integration across Different Psychedelic Compounds

To investigate the neural circuits underlying perceptual integration and how they are modulated by psychedelics, we conducted whole-brain c-Fos mapping in mice following task performance and homecage conditions after treatment with psilocybin (1 mg/kg), ketamine (30 mg/kg), or saline. Brain tissue was collected ∼110 minutes after the start of the task, and we quantified c-Fos expression across dozens of regions using an automated pipeline (Figure 3A–B)^18,19^. This approach enabled us to identify brain regions exhibiting increased activity associated with sensory evidence accumulation and decision-making across different drug conditions.

**Figure 3.**
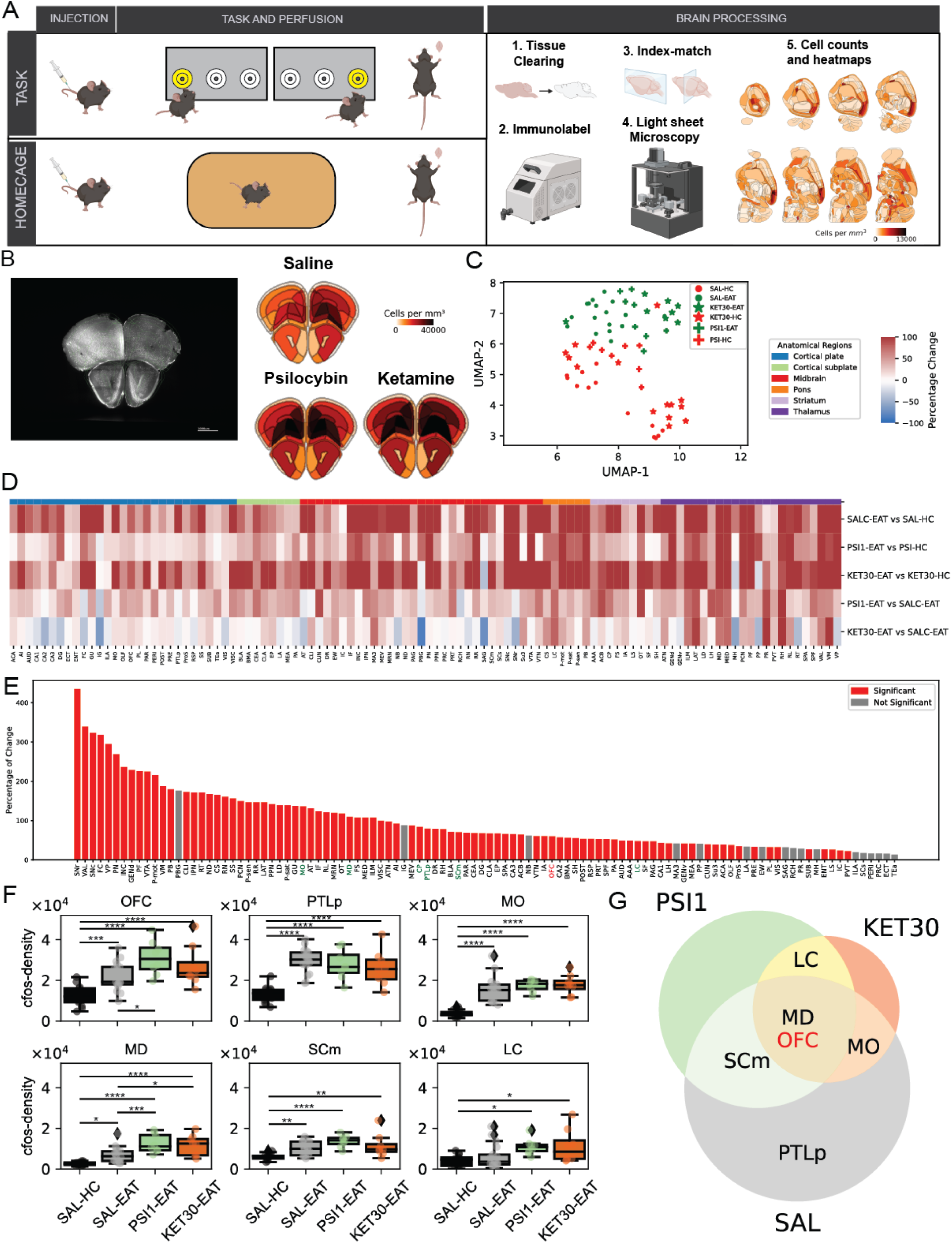
Whole-brain c-Fos mapping reveals a conserved neural circuit for perceptual evidence accumulation modulated by psychedelics. **(A)** Experimental design. Mice were injected with saline, psilocybin (1 mg/kg), or ketamine (30 mg/kg), then either performed the free-response perceptual accumulation task or remained in the homecage. Brains were processed at LifeCanvas via iDISCO+ tissue clearing, immunolabeling, and whole-brain light-sheet microscopy to quantify c-Fos expression. **(B)** Example coronal images showing c-Fos expression in task-performing animals across all treatment groups, during the task performance. **(C)** UMAP embedding of whole-brain c-Fos expression patterns. Task-engaged animals (EAT) cluster separately from homecage controls (HC), independent of drug condition. Psilocybin and ketamine also produce distinct activity patterns relative to saline during task performance. **(D)** Heatmap showing normalized c-Fos expression across all brain regions and conditions. Most regions exhibit increased activity during task engagement. **(E)** Bar plot ranking brain areas by the number of significant task-related c-Fos increases across conditions. Red bars indicate regions with significant upregulation across multiple comparisons. **(F)** c-Fos expression levels in six key brain regions involved in evidence accumulation: orbitofrontal cortex (OFC), posterior parietal cortex (PTLp), motor cortex (MO), mediodorsal thalamus (MD), superior colliculus (SCm), and locus coeruleus (LC). Both task and psychedelics increase c-Fos expression in these areas during task engagement compared to controls. **(G)** Venn diagram showing the overlap of task- and drug-responsive regions. OFC and MD are commonly activated by both psychedelics. SCm, MO, and LC are selectively engaged by either psilocybin or ketamine, while PTLp is primarily engaged under all conditions. Notations: * = p<0.05, ** = p<0.01, *** = p<0.005, **** = p<0.001.

Unsupervised clustering and UMAP projections of c-Fos expression revealed a clear separation between mice that performed the task and those in the homecage condition, independent of treatment. This distinction indicates a consistent neural signature associated with perceptual integration. Additionally, among task-performing animals, psilocybin, ketamine, and saline-treated groups exhibited distinct clustering patterns, suggesting that each drug induces unique yet overlapping neural activation profiles during decision-making (Figure 3C).

To further characterize these differences, we compared the percentage change in c-Fos expression across brain regions relative to their homecage controls, identifying both shared and drug-specific activation patterns. Many cortical, subcortical, midbrain, and thalamic regions exhibited significantly higher c-Fos expression in task-performing mice compared to homecage controls, reinforcing the broad neural engagement required for perceptual decision-making. Additionally, psilocybin and ketamine induced distinct activation patterns compared to saline, indicating their specific influence on neural circuits involved in perceptual integration (Figure 3D-E). These compound-specific patterns were further reflected in large-scale functional connectivity and network topology, as visualized by correlation matrices and graph representations (Supplementary Figure 4A).

Overall, the majority of brain regions demonstrated a significant increase in c-Fos expression during task performance (Figure 3E). Notably, several cortical and subcortical areas exhibited significant activation differences, with multiple regions previously implicated in perceptual integration emerging as key components of this circuit. Among the most activated regions, we identified the orbitofrontal cortex (OFC; *F = 12.74, p < 0.0005*), medial dorsal thalamus (MD; *F = 16.52, p < 0.0005*), superior colliculus (SCm; *F = 9.14, p < 0.0005*), locus coeruleus (LC; *F = 4.23, p = 0.011*), posterior parietal cortex (PTLp; *F = 18.02, p < 0.0005*), and motor cortex (MO; *F = 25.24, p < 0.0005*) (Figure 3E-F). These findings suggest that psychedelics engage a recurrent cortical-subcortical network involved in sensory evidence integration and decision-making, providing further insight into their role in modulating cognitive processing.

Finally, a Venn diagram analysis comparing brain regions significantly activated during task performance (relative to homecage) in the saline condition with those further modulated by psilocybin or ketamine (relative to saline during the task) revealed key areas involved in perceptual integration and psychedelic-induced neural modulation. The orbitofrontal cortex (OFC) and medial dorsal thalamus (MD) were consistently engaged during task performance and further enhanced by psychedelics. Additionally, the LC was uniquely engaged under psilocybin and ketamine, whereas the SCm showed stronger activation with psilocybin, and the MO was more activated with ketamine, while the PTLp remained similarly active across conditions (Figure 3F, Supplementary Figure 4B).

### Psychedelics Reshape OFC Decision-Related Dynamics by Modulating Signal Strength, Temporal Structure, and Population Decorrelation

To investigate how psychedelics alter decision-related computations in cortical circuits involved in perceptual integration, we focused on the orbitofrontal cortex (OFC)—a region consistently modulated by the task and psychedelics in our whole-brain mapping and previously implicated in perceptual decision-making^20–22^. While OFC has been examined in decision-making contexts, its single-neuron population dynamics during free-response evidence accumulation—and how these dynamics are reorganized by psychedelics—have not been previously characterized. We recorded 2,777 OFC neurons using cellular-resolution calcium imaging as mice performed the task (Figure. 4A–B). We employed logistic regression to predict behavioral variables from neural activity^23^. The model accurately decoded choice, reward, and decision time with over 75% accuracy, confirming robust task-related representations in OFC (>75% accuracy; Figure 4C–D), with neurons classified into task-selective subgroups based on the sign and magnitude of their beta weights (Supplementary Figure 5).

**Figure 4.**
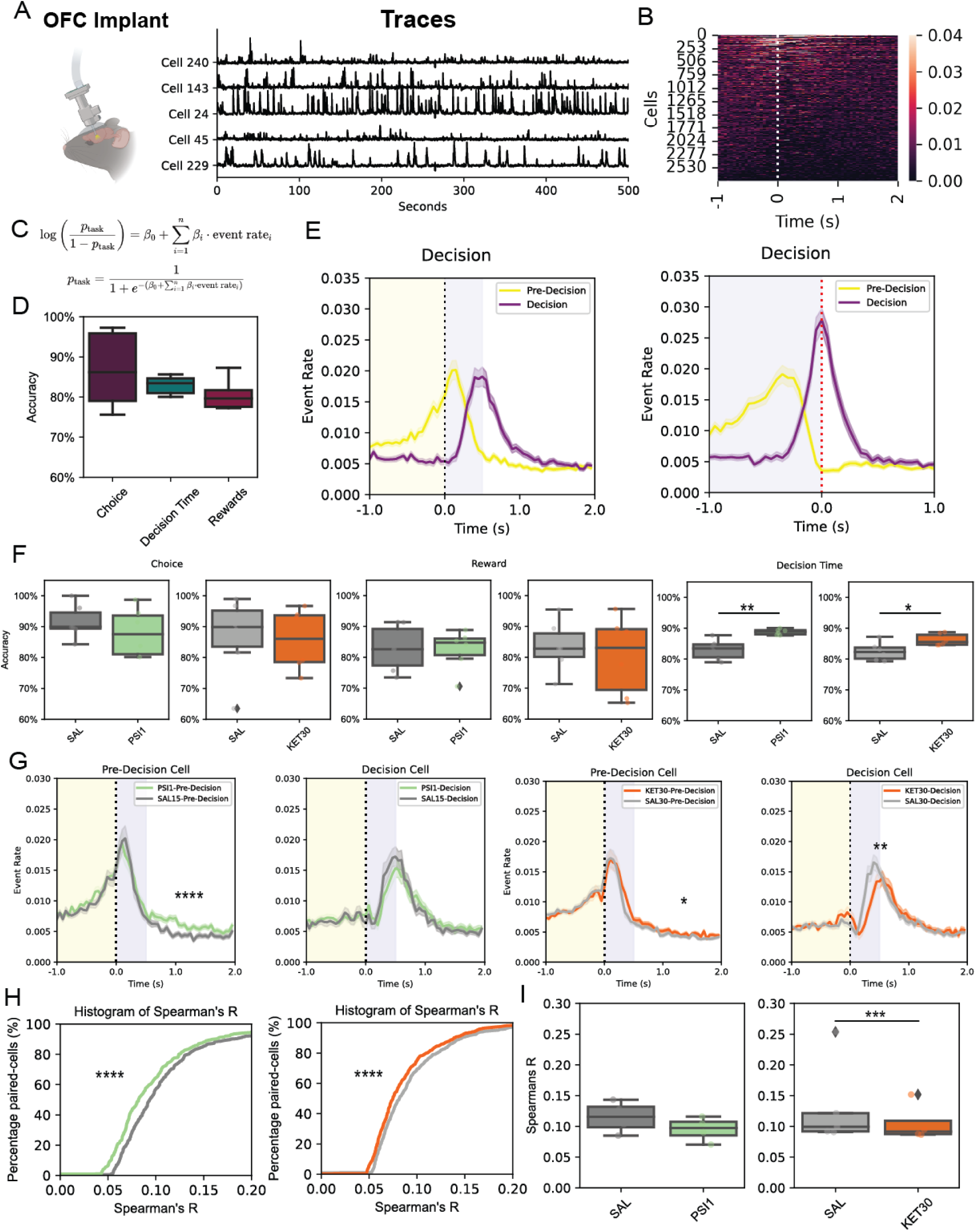
Psychedelics reshape OFC decision-related dynamics by modulating signal strength, decodability and temporal structure of decision time related cells. **(A)** Schematic of one-photon calcium imaging setup during behavior, with example traces from OFC neurons recorded through GRIN lenses. **(B)** Heatmap showing trial-aligned event rates for individual decision-selective OFC neurons, centered around the time of choice (red dashed line). **(C)** Logistic regression model used to classify neurons based on their ability to predict behavioral outcomes (choice direction, reward, and decision time) from event rates. **(D)** Classification accuracy of logistic models. OFC activity could reliably predict choice direction, reward outcome, and decision timing with >75% accuracy. **(E)** Average calcium event rate of decision-selective neurons (n = 476 neurons) aligned to trial initiation (left) and choice (right). Pre-Decision cells (n = 267 neurons) ramped up before the choice and Decision cells (n = 209 neurons) peaked during choice. **(F)** Accuracy of logistic regression models trained on OFC calcium event rates to predict behavioral variables. Psilocybin and ketamine significantly improved model accuracy for predicting decision time, with modest effects on choice and reward prediction. **(G)** Trial-aligned event rate of pre-decision and decision-selective cells under psilocybin (green, left panels; Both: n = 551 neurons, Pre-Decision: n = 317 neurons, Decision: n = 194 neurons) and ketamine (orange, right panels; Both: n = 601 neurons, Pre-Decision: n = 395 neurons, Decision: n = 206 neurons), compared to saline (Both: n = 557 neurons, Pre-Decision: n = 331 neurons, Decision: n = 226 neurons). Both drugs significantly delayed and reduced peak activity in decision cells, consistent with slower decision times. Psilocybin significantly decreased post-decision activity; ketamine induced a smaller but significant reduction in both post-decision and decision cell activity. **(H)** Cumulative histograms of Spearman’s correlation coefficients for temporal structure in decision cell activity. Both psilocybin and ketamine shifted distributions toward lower correlations. **(I)** Per-animal averages of Spearman’s R. Ketamine significantly reduced temporal consistency, indicating disrupted temporal structure in decision-related activity. Notations: * = p<0.05, ** = p<0.01, *** = p<0.005, **** = p<0.001.

Focusing on decision time-selective neurons, we aligned their activity to trial events to resolve their temporal structure. Focusing on decision time-selective neurons, we aligned their activity to trial events and found that these cells formed a temporal sequence: Pre-decision cells ramped prior to commitment, while decision cells peaked sharply at the moment of choice (Figure 4E, Supplementary Fig. 5). To assess how psychedelics affect these dynamics, we retrained models under drug conditions. Both psilocybin and ketamine significantly improved decision time decoding, with no such effect under cocaine (*Psilocybin: T = 4.73, p = 0.005; Ketamine: T = 3.10, p = 0.027*; Figure 4F, Supplementary Fig. 3I). This enhancement suggests that psychedelics increase the temporal precision of decision-related coding. Trial-aligned averages revealed delayed and reduced decision cell peaks under both compounds (Figure 4G), mirroring the longer decision times observed behaviorally. This finding aligns with the hypothesis that psychedelics alter the temporal structure and precision of decision-related signals, potentially making commitment points more identifiable at the circuit level^24,25^. Psilocybin and ketamine also elevated post-trial activity in pre-decision cells. Furthermore, both drugs altered choice- and reward-selective dynamics, pointing to widespread reorganization of task-relevant activity in OFC (Supplementary Figure 6).

To assess the coordination of neural activity during decision-making, we computed pairwise Spearman correlations between decision-selective OFC neurons, based on their activity patterns across trials. Both psilocybin *(Mann-Whitney U; p < 0.0005; Bootstrapped mean difference: 0.013 and confidence intervals: [0.0054, 0.021]*) and ketamine (*Mann-Whitney U; p < 0.0005; Bootstrapped mean difference: 0.006 and confidence intervals: [0.0012, 0.0105]; T = 27.95, p = 0.001*) shifted the distribution of correlations toward lower values (Figure 4H-I). These reductions in inter-neuronal correlation reflect a decorrelation of ensemble activity, a phenomenon known to enhance population coding by increasing representational dimensionality and reducing shared noise^24,25^. At the network level, such decorrelation has been linked to increased information processing capacity, state flexibility, and cognitive adaptability—features that align with the proposed neural signatures of psychedelic brain states, including heightened signal complexity and functional reorganization^26,27^.

In summary, psychedelics reshape OFC dynamics by delaying commitment signals, enhancing decision time decoding, and decorrelating population activity. These effects increase the separability of decision states and likely support the transition to a more deliberate, flexible inference strategy. The convergence of these neural changes with behavioral effects observed under psilocybin and ketamine provides circuit-level support for the REBUS model, which posits a weakening of top-down priors and enhancement of bottom-up evidence processing^3^.

## Discussion

Our results show that psychedelics fundamentally alter perceptual decisions by modulating the interplay between prior expectations and sensory evidence. The shift we observed—slower decisions paired with improved accuracy—reflects a reconfiguration of decision policy that favors real-time evidence accumulation over fast, bias-driven responding. Drift diffusion modeling revealed that psychedelics increased decision thresholds without altering sensory gain, consistent with reduced prior precision and greater demand for evidence before commitment. This strategy—slower but more accurate—suggests a shift in decision-making policy, prioritizing cautious, sensory-guided inference over speed or habit. Such a shift may reflect a reweighted internal cost function, where the cognitive or behavioral cost of acting on biases increases relative to the value of accumulating evidence, making prior-driven shortcuts less favorable under psychedelics. This is precisely the behavioral signature expected when high-level priors are relaxed and bottom-up input gains greater influence in shaping belief updating^3^. GLM-HMM modeling further showed that these behavioral changes reflected transitions from prior-dominated to sensory-engaged cognitive states, characterized by reduced influence of past choices and increased responsiveness to real-time input. These transitions were robust, dose-dependent, and compound-specific, indicating a convergent computational outcome across distinct receptor targets. Therefore, rather than enhancing perceptual acuity directly, psychedelics appear to relax the weight of prior beliefs, effectively raising the threshold for commitment and prolonging deliberation. These results align with the core prediction of the REBUS model^3^: that psychedelics relax the precision of high-level priors, thereby flattening hierarchical predictive coding and opening the system to bottom-up information. Human studies support this mechanism: psychedelics reduce mismatch negativity^28^ and weaken top-down control from associative networks, such as the default mode network, over sensory areas^29–31^. These convergent findings across species suggest a conserved computational signature of psychedelic action—namely, the relaxation of high-level priors and enhanced sensitivity to bottom-up input—that may underlie both the acute perceptual effects and the therapeutic potential of these compounds.

Whole-brain c-Fos mapping revealed a conserved decision-making network encompassing cortical and subcortical regions—including the orbitofrontal cortex (OFC), medial dorsal thalamus (MD), superior colliculus (SCm), locus coeruleus (LC), motor cortex (MO), and posterior parietal cortex (PTLp)—recruited across both saline and psychedelic conditions. Compound-specific changes across these nodes further suggest that psychedelics reconfigure, rather than disrupt, the distributed circuitry supporting perceptual decisions—tuning the balance between top-down and bottom-up processing to enable more flexible and context-sensitive inference^4,32^. Among these, the OFC emerged as a key hub, consistently activated across all treatment groups and strongly modulated by both psilocybin and ketamine. Given its central role in belief updating, value-guided behavior, and encoding task structure^11,12,33^, the OFC is a likely site of convergence for priors and sensory evidence, making it particularly relevant for testing the REBUS model. Our in vivo calcium imaging further revealed that psychedelics preserved decision-related selectivity in OFC neurons while reshaping their temporal dynamics. Psilocybin and ketamine delayed and dampened peak decision activity, consistent with prolonged evidence accumulation. At the ensemble level, both compounds induced significant decorrelation among decision-selective neurons—a transformation linked to increased dimensionality and cognitive flexibility^34,35^. These changes mirror theoretical accounts in which psychedelics elevate cortical entropy and reduce the precision of top-down priors^3,36^. Notably, these population-level reconfigurations align with previous studies demonstrating that psilocybin and ketamine reorganize prefrontal functional connectivity and promote neural plasticity in frontal circuits, including OFC^37,38^. Crucially, the observed circuit-level reconfiguration converged with behavioral shifts toward longer response times and improved accuracy, providing mechanistic evidence that psychedelics relax priors to support more deliberate, sensory-driven inference.

Several limitations warrant discussion. First, while we identified a conserved circuit for evidence accumulation and mapped its modulation by psychedelics, the causal contributions of individual regions remain unresolved. Second, our task captures perceptual decision-making in a constrained sensory context; whether similar neural dynamics and state transitions emerge during more abstract or socially embedded forms of inference remains an open question. Third, although psilocybin and ketamine target distinct molecular pathways, their convergence at the circuit and computational levels suggests a shared functional outcome. Expanding this work to include a broader panel of serotonergic, glutamatergic, and mixed-mechanism psychedelics will help distinguish general principles from compound-specific effects.

Psychedelics restructure perceptual decision-making by shifting behavior from rapid, prior-biased strategies toward slower, evidence-guided inference. Through integrated behavioral modeling and circuit-level analysis, we show that psilocybin and ketamine attenuate prior-driven biases, elevate decision thresholds, and reorganize orbitofrontal dynamics. These findings establish a mechanistic framework by which psychedelics relax top-down constraints, supporting core predictions of the REBUS model in a tractable animal system.

## Acknowledgments

We thank Julie Gomez, Arula Ratnakar, Zeynep Ozturk, Christa Rose, Lorenzo Gatti, Cameron Chieppo and Sina Analoui for help with collection of rodent data and Albit Caban for his help with the GRIN lens surgeries and recordings. We thank Josh Sanders for feedback on the behavioral training software. This work was supported by NIH Transformative Award, the Ludwig Family Foundation, the Air Force Office of Scientific Research award (FA9550-21-1-0310), the Pew Scholars Program in the Biomedical Sciences, the Chan-Zuckerberg Initiative, and Neurophotonics Center at Boston University and the Center for Systems Neuroscience distinguished fellowship award to CDS.

## Contributions

CDS designed the experiments. CDS, SAA, AKL and BSL performed the experiments. CDS and RS analyzed the data. CDS, BBS and SR wrote the manuscript. All authors contributed to the conceptualization and execution of the study and edited the manuscript.

## Materials and methods

### Rodents

All experiments and procedures were approved by the Boston University Animal Care and Use Committee. C57BL/6NJ adult mice (n = 111, ages 2–8 months) and a Thy1-GCaMP6s transgenic C57BL/6J line (GP4.3Dkim/J, n = 6, ages 2–8 months) were obtained from The Jackson Laboratory. Both male and female mice were used, and they were trained concurrently in the same room but in separate operant chambers.

Mice were water-restricted to maintain 80–90% of their baseline body weight and received at least 1 mL of water per day. Each reward consisted of 5 μL of 10% sucrose solution (100 g/L), delivered consistently in each trial. Mice were housed on a 14:10 h light:dark cycle, with lights on during local daytime (Boston, MA, USA). Animals were housed in groups of 2–4 per cage. Each morning, mice were transported to the training room for behavioral sessions. Training sessions lasted 1 hour, 5 days per week, occurring either in the late morning (10 am–12 pm) or early afternoon (12 pm–3 pm).

### Rodent behavioral control system

We modeled the behavioral apparatus after previously described designs^39,40^. Mice were trained in custom acrylic chambers containing three nose-poke ports. The ports (Sanworks or custom-made) were each outfitted with a visible LED for stimulus presentation, a peristaltic pump for liquid reward delivery, and an infrared beam-break sensor (emitter and photodetector) to detect nose pokes. The chamber floors were lined with bedding.

Task control software was written in MATLAB and interfaced with a Teensy-based microcontroller running the Bpod framework (Sanworks). A custom Python application coordinated multiple Bpod devices from a single computer, using a modified version of the Bpod MATLAB library (edited Bpod library: https://github.com/RatAcad/Bpod_Gen2; Custom python application: https://github.com/RatAcad/BpodAcademy). Further details of the hardware and software design are available in the referenced repositories.

### Rodent training pipeline

Training took 3–4 weeks per mouse and progressed through three stages. In the first stage (1–3 days), mice were rewarded for inserting their nose into a side port with an illuminated LED (400 trials per session). In the second stage, mice had to nose-poke the center port and then a side port with the LED illuminated to receive reward (400 trials). In the third stage, mice were required to nose-poke the center port and then a side port with a flashing LED for reward.

Once mice reached a performance criterion of 90–100% correct choices over 400 trials in the third stage, we increased task difficulty by introducing flashes on the incorrect side. The probability of an incorrect-side flash was gradually raised from 0% to 20% (in steps: 100:0, then 90:10, then 80:20 correct:incorrect flash ratio). Mice trained for 1–5 days at each probability condition. After completing all training stages and conditions, mice performed the final task with an 80:20 flash probability ratio^9^.

### Rodent behavioral task

During task sessions, mice could move freely within the chamber. At the start of each trial, the center port LED turned on, indicating the mouse could initiate the trial by nose-poking that port. Upon a nose poke in the center port, a single flash was simultaneously presented at both left and right ports to signal trial onset. Thereafter, flashes were generated according to a Bernoulli process: every 100 ms a flash appeared in either the left or right port, with an 80% probability of occurring on the designated “correct” side (and 20% on the opposite side). The correct side for each trial was chosen randomly. The sequence of flashes continued until the mouse made a choice by nose-poking one of the side ports, upon which the stimulus stream immediately ceased. If the mouse chose the correct side, the corresponding port’s LED remained illuminated for 3 s and a 5 μL sucrose reward was delivered. If the mouse chose incorrectly, all lights turned off for a 5 s timeout with no reward, and the next trial became available after a 2 s delay. Each trial had a maximum duration of 8 s; if no response was made by 8 s, the trial was counted as an omission.

### Drug administration protocols

All drugs were administered via intraperitoneal (i.p.) injection once per week to allow full washout and minimize carryover effects. Each animal received only one type of compound—psilocybin (USONA Institute, Madison, WI, USA; n = 42), ketamine (Ketamine Hydrochloride Injection, 100 mg/mL; Covetrus; SKU; n = 30), or cocaine (Cocaine Hydrochloride; Sigma-Aldrich, Cat. No. C5776; n = 12)—but was exposed to multiple doses of that compound in separate sessions using a within-subjects design. Psilocybin was administered at 0.5, 1, and 3 mg/kg; ketamine at 10, 30, 50, and 70 mg/kg; and cocaine at 5, 10 and 20 mg/kg. All drugs were dissolved in 0.9% sterile saline and injected at a volume of 10 mL/kg. Animals received saline injections in control days. Behavioral testing commenced 10 minutes after injection for psilocybin, 30 minutes after ketamine injections, and 5 minutes after injection for cocaine, in line with previous pharmacokinetic studies indicating peak central effects within these time frames. The order of doses was counterbalanced across animals within each drug group to control for potential order effects.

### Data analyses

Data analysis was conducted using Python version 3.7.1 and R version 4.3.1. Trial events were aligned via TTL pulses recorded from the behavioral control system, enabling synchronization of trial initiation, stimulus presentation, and response timestamps across systems. Each animal contributed multiple behavioral sessions per pharmacological condition. Trial-level data were aggregated across sessions within animal and condition unless otherwise specified.

The total number of trials per session was computed for each animal across all sessions. Mice during baseline condition (before any treatment was administered) completed an average of 296 ± 104 trials per session (n = 743 sessions). During saline administration they completed 289 ± 105 trials per session (n = 557 sessions). For psilocybin, trial counts per session ranged from 186 to 299 depending on dose (0.5–3 mg/kg). Ketamine sessions ranged from 166 to 208 trials per session across the 10–70 mg/kg range. Cocaine-treated animals exhibited the highest throughput, averaging ∼390 trials per session. These per-session estimates were used to normalize behavior and imaging-derived metrics where applicable.

Accuracy was calculated as the number of correct trials divided by the total number of completed trials and expressed as a percentage. Response time (RT) was defined as the interval between the onset of the first flash and the animal’s side-port response. These behavioral metrics—accuracy and RT—were calculated over time using a 5-minute rolling average across the 60-minute session. To assess task engagement dynamics, we also quantified the trial rate per session by averaging the number of trials completed per minute across sessions (Supplementary Figure 2).

To model the relationship between RT and accuracy (Figure 2C), we applied a generalized additive mixed model (GAMM) using the mgcv package in R^41^. Accuracy was averaged in 0.2-second RT bins over a 2-second window, and only bins with more than 50 trials were retained for analysis. Each condition was modeled independently across pooled trials.

Statistical comparisons, including one-way ANOVAs, independent and paired t-tests, and Pearson or Spearman correlations, were conducted using the pingouin package in Python^42^. Where appropriate, p-values were corrected for multiple comparisons using Bonferroni corrections.

### Simulation Analysis

To benchmark behavioral performance, we simulated a perfect accumulator model under conditions that matched the task’s generative structure (Figure 2B). Simulations included 10,000 trials for each fixed RT value ranging from 100 ms to 2 s in 200 ms increments. On each trial, a generative flash probability of 80% was assigned to one side (left or right), determined randomly via a coin flip. Individual flashes were generated independently using a Bernoulli process, and sequences terminated once the simulated RT was reached.

The perfect accumulator counted the total number of flashes per side and always selected the side with more flashes. If both sides had equal flash counts at the decision time, the model selected a side at random. For each RT condition, the simulation yielded an estimate of expected accuracy and reward rate under ideal evidence integration. These outputs served as theoretical upper bounds for behavioral performance and informed interpretation of the observed speed-accuracy trade-offs.

### GLM and GLM-HMM

To quantify the contribution of sensory evidence and choice history to behavior, we fit generalized linear models (GLMs) with a logit link function predicting rightward choice probability. Regressors included (i) normalized pulse difference between sides, (ii) win-stay/lose-switch strategy, and (iii) previous choice. The intercept captured side bias.

Analyses were restricted to trials from the 80% vs. 20% flash distribution. GLMs were implemented using standard maximum likelihood estimation methods^43,44^.

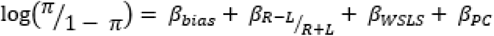

Model fits were evaluated using log-likelihood per trial expressed in bits, defined as log-likelihood divided by 𝑛 * 𝑙𝑜𝑔(2), following Ashwood et al. (2022). To capture non-stationary strategies, we used a hidden Markov variant (GLM-HMM), in which each latent state corresponded to a distinct GLM with state-specific weights and transition probabilities. Models were fit using expectation-maximization with hyperparameters σ = 2 and α = 2. The hyperparameters dictate the strength of the prior on the beta coefficients of the GLM and transition matrix respectively. The sigma parameter is the variance of a zero-mean Gaussian prior placed over the GLM weights; higher values of sigma correspond to a wider Gaussian allowing larger values of weights and thus a less informative prior. The α parameter is the value for a symmetrical dirichlet distribution that is the conjugate prior for each row of the transition matrix of the HMM, A. A value of 1 is an uninformative or flat prior, whereas higher values of α cause the transition matrix to be more homogenous.This approach extends previous work applying latent-state models to decision-making behavior^13,45^. For comparison, a two-state lapse model was also fit, capturing alternations between engaged (stimulus-dependent) and lapse (stimulus-independent) behavior.

### Drift diffusion models

To infer latent decision dynamics, we applied drift diffusion models (DDMs), which describe decisions as the result of noisy evidence accumulation between two boundaries^14,15,46^. Decision-making was governed by drift rate (v), boundary separation (a), starting point (z), and non-decision time (ndt). Drift rate reflects the speed of accumulation, boundary separation the evidence required to commit, starting point the initial bias, and non-decision time accounts for sensory and motor delays^9^ . Evidence was modeled as:

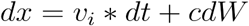

where 𝑑𝑥_𝑖_(𝑡) is the evidence change on trial 𝑖, 𝑐 is a noise constant, and 𝑑𝑊_𝑖_ is Gaussian noise. Trial-to-trial variability in v, z, and ndt was modeled as Gaussian, uniform, and half-normal distributions, respectively. Models were fitted using a custom R implementation (rddm; https://github.com/gkane26/rddm) by minimizing the χ² distance between simulated and empirical RT histograms. Parameters were optimized via differential evolution to match both accuracy and RT distributions.

### State-Resolved DDM via GLM-HMM Posterior Inference

To examine whether latent behavioral states correspond to distinct accumulation dynamics, we used posterior state probabilities from the GLM-HMM to classify trials into latent states. Trials with posterior probabilities ≥ 0.8 were assigned to their most likely state. Separate DDMs were then fit to each state’s trial subset per animal.

This approach enabled estimation of state-specific parameters, revealing whether sensory-engaged states were associated with higher drift rates and decision thresholds compared to biased or lapse states. Models were fit and compared using the same χ²-based procedure. This joint modeling framework allowed us to link latent strategy transitions with underlying decision computations, providing a multi-scale mechanistic account of how psychedelics alter inference and evidence accumulation.

### Whole-brain c-Fos staining and imaging

To investigate brain-wide activity associated with perceptual decision-making under psychedelic or saline conditions, we quantified c-Fos expression using whole-brain light-sheet microscopy. Animals (C57BL/6NJ) were randomly assigned to receive saline (HC: n = 14, EAT: n = 15), psilocybin (1 mg/kg; HC: n = 8, EAT: n = 8), ketamine (30 mg/kg; HC: n = 15, EAT: n = 8), or cocaine (20 mg/kg; EAT: n = 8) and were either placed in the behavioral task or maintained in their homecage. For task-performing animals, injections occurred 10–30 minutes (1 minute for cocaine) before the start of the session, and mice were allowed to complete trials freely in the operant chamber for a fixed duration of 60 minutes. This timing was chosen to align the peak of drug-evoked gene expression with robust task engagement while avoiding confounds from late-session fatigue or satiety.

Approximately 110 minutes after the onset of the behavioral session, animals were deeply anesthetized with isoflurane and transcardially perfused with cold phosphate-buffered saline (PBS), followed by ice-cold 4% paraformaldehyde (PFA) in PBS. Brains were extracted carefully, post-fixed in 4% PFA overnight at 4 °C, and then transferred to PBS for shipment.

Fixed tissue samples were submitted to LifeCanvas Technologies for whole-brain processing, immunolabeling, and imaging^19,47^. Brains were first treated with the SHIELD tissue preservation protocol to maintain structural and molecular integrity while enabling high-throughput processing^47^. Delipidation was performed using the SmartBatch+ active clearing system to promote uniform antibody penetration across tissue volume^47,48^. Samples were then incubated with rabbit anti-c-Fos primary antibody (Abcam, ab214672) followed by species-matched fluorescent secondary antibodies. After immunostaining, brains were index-matched with EasyIndex solution and imaged using the SmartSPIM light-sheet microscope to acquire high-resolution, whole-brain fluorescence volumes^19^.

Imaging datasets were registered to the Allen Mouse Brain Atlas using SmartAnalytics software, and c-Fos+ cells were automatically detected and quantified across more than 150 anatomically defined regions. Differential expression analyses were performed by comparing region-specific c-Fos densities across conditions (drug vs. saline; task vs. homecage), yielding brain-wide maps of activation associated with task performance and psychedelic modulation.

### Functional network construction from c-Fos expression

To characterize large-scale co-activation patterns across brain regions, we constructed functional c-Fos networks by correlating regional c-Fos expression levels across animals within each condition (Supplementary Figure 4). For each experimental group, we calculated Spearman correlation coefficients between all pairs of brain regions (n = 150, as defined by the Allen Brain Atlas), using the density c-Fos+ cell counts for each region per animal. This yielded symmetric correlation matrices reflecting the degree of covariation in activity between regions. Networks were visualized by thresholding the correlation matrices at a fixed sparsity level (top 10–15% of edges) to retain the strongest positive correlations. Graph theoretical measures—including node strength, modularity, and participation coefficient—were computed using the Brain Connectivity Toolbox to quantify network organization and integration across conditions. Network diagrams and spatial projections were rendered using custom Python scripts^49^. These analyses enabled comparison of how psychedelic compounds differentially engage and reorganize mesoscale functional architecture during task performance.

### GRIN lens implantation for in vivo calcium imaging

To enable cellular-resolution calcium imaging in the orbitofrontal cortex (OFC), we implanted gradient refractive index (GRIN) lenses (0.5 mm diameter; Inscopix) in Thy1-GCaMP6s transgenic mice (n = 6). Mice were anesthetized with isoflurane (4-5% induction, 1–2% maintenance in oxygen) and placed in a stereotaxic apparatus. Local anesthesia (lidocaine, 0.5%, topical) was administered. The skull was exposed, and a small craniotomy was made above the OFC using stereotaxic coordinates: anteroposterior (AP) +2.4 mm, mediolateral (ML) ±1.0 mm from bregma. After carefully removing the dura, a stainless steel insertion cannula with a tapered tip was slowly advanced to –2.1 mm to create a tract for lens implantation.

The GRIN lens was slowly lowered to a final depth of dorsoventral (DV) –2.4 mm relative to the skull surface, ensuring minimal tissue compression. The lens was secured to the skull using light-curable dental cement (Metabond) and affixed to a custom titanium baseplate for future attachment of the miniature microscope. The incision was closed around the implant base with tissue adhesive. Postoperative care included daily monitoring and analgesia (meloxicam, 5 mg/kg, and buprenorphine, 0.1 mg/kg s.c.) for 3 days. Mice were allowed to recover for at least 3 weeks before beginning imaging sessions to allow tissue stabilization and lens clearance.

### Calcium imaging preprocessing and event rate extraction

GCaMP6s-expressing mice implanted with GRIN lenses targeting the orbitofrontal cortex (OFC) were imaged during task performance using a miniature fluorescence microscope (nVista 3.0, Inscopix). Recordings were performed at 20 Hz using the Inscopix Data Acquisition Software (IDAS), and TTL pulses sent from the BPOD behavioral control system were logged to ensure precise alignment between imaging frames and behavioral events. A hardware trigger from the behavioral controller was used to initiate video recording, and all TTL events (e.g., trial start, choice, reward) were simultaneously recorded as timestamped annotations within the video metadata.

Raw videos were spatially downsampled by a factor of 2 and motion-corrected using the Inscopix Data Processing Software (IDPS, v1.7.1). After motion correction, the ΔF/F signal was computed for each pixel as ΔF/F = (F – F₀)/F₀, where F₀ was defined as the 8th percentile of the fluorescence signal over a 60-second sliding window. Spatial components corresponding to individual neurons were extracted using CNMF-E (Constrained Non-negative Matrix Factorization for Endoscopic data) implemented via the open-source CaImAn package in Python.

The inferred calcium traces were deconvolved using an autoregressive (AR-1) model to estimate the underlying spike probability. To obtain binarized event rasters, we applied an adaptive threshold to the deconvolved signal (4 standard deviations above baseline noise and 0.2 seconds for the smallest decay time), defining a calcium event as any frame in which the signal exceeded this threshold. These binarized events were then aligned to behavioral timestamps using the recorded TTL pulses, allowing quantification of trial-by-trial event rates for each neuron in relation to specific task epochs (e.g., stimulus onset, decision, outcome). Event rasters and peri-event time histograms (PETHs) were constructed for each cell, time-locked to behavioral markers. Trial-averaged activity was normalized across conditions and used for regression-based cell classification.

### Neuronal classification and statistical comparison

To identify neurons encoding decision-relevant variables, we implemented a logistic regression framework in which binarized calcium event rates were used to predict trial-by-trial behavioral outcomes. For each neuron, a generalized linear model (GLM) with a binomial distribution and logit link function was fit to one of three behavioral features: (1) choice direction (left vs. right), (2) reward outcome (correct vs. error), or (3) decision time (binned into flashes).

For the choice and reward models, calcium events were binned into 50 ms windows and aligned to trial initiation or reward delivery, respectively. The sum of binarized events across a 1-second window following each alignment point was used as the predictor variable. For decision time, calcium events were binned into 100 ms windows aligned to each individual flash onset, starting from trial initiation and continuing through to the moment of choice. A value of 1 was appended to the final bin corresponding to the time of choice to signal commitment, while bins with no events retained a value of 0. The sum of binarized events for each flash-aligned bin served as the predictor set for modeling decision latency.

Each model took the form:

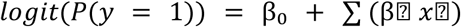

where y is the behavioral label, xt is the event rate in time bin t, and βt are the learned weights. Prediction accuracy was evaluated using 5-fold cross-validation.

Neurons were classified as selective if they had non-zero weights after L1 penalization, indicating predictive contribution and leveraging its ability to both reduce overfitting and perform feature selection by shrinking irrelevant coefficients to zero^23^. The sign of each non-zero β coefficient indicated response polarity. Model fitting used sklearn.linear_model.LogisticRegression with penalty=’l1’ and solver=’liblinear’, optimizing the logistic loss with an added L1 penalty term:

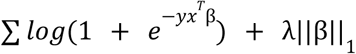

This promotes sparse solutions by selecting only informative neurons. By default, sklearn uses 5-fold cross-validation to select the best value of the hyperparameter λ.

### Non-parametric permutation testing for group comparisons

To assess statistical differences in calcium event rates across experimental conditions, we implemented a custom non-parametric permutation test based on shuffled group labels. Binarized calcium activity (0/1 events) was first segmented into time windows of interest: a primary test window, and flanking pre- and post-decision windows. For each neuron type (e.g., choice, reward, decision) and trial type (e.g., correct, error), we extracted calcium event rates from the selected time window at a sampling rate of 20 Hz (downsampled to 0.05 s bins).

For each condition pair (e.g., saline vs. psilocybin), we computed the observed difference in mean firing rate across neurons. To evaluate the significance of this difference, we generated a null distribution by pooling all trials across both conditions and randomly permuting group labels (n = 1,000 permutations). For each permutation, we recalculated the mean event rate difference. This process was repeated for the pre-, test-, and post-decision windows.

An empirical two-tailed p-value was computed as the proportion of permuted differences greater than or equal to the observed difference in absolute value:

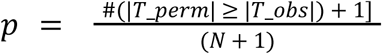

where Tobs is the observed difference in mean firing rate, and T_perm represents the permuted differences across N=1000 iterations. This analysis was repeated across all trial types and neuron classes, producing a set of window-specific p-values for each pairwise condition comparison.

### Correlation analyses of decision-related neurons

To examine the temporal coordination and trial-to-trial variability of decision-selective neurons, we analyzed binarized calcium activity aligned to behavior. Only neurons previously identified as decision cells via L1-regularized logistic regression were included. For each animal and condition, activity traces from selected cells were aligned to trial initiation and binned into non-overlapping windows (50 ms) across the full trial duration. A binary raster was generated for each trial, and summed activity within each bin was used to estimate firing rate.

To assess how neuronal activity covaried across trials, we calculated Spearman rank correlations between all pairs of decision cells using the concatenated activity matrix across trials. Correlations were computed for the full trial window. The lower triangle of the resulting correlation matrices was extracted for analysis, and statistical significance was assessed via permutation-based p-values. To compare the distributions of Spearman’s rank correlation coefficients across experimental conditions with unequal sample sizes, we employed a non-parametric bootstrapping approach. For each condition pair, cells with significant correlations (p < 0.05) were selected, and Spearman’s R values were extracted. To compare Spearman’s correlation distributions across conditions with unequal sample sizes, we used the Mann–Whitney U test, a non-parametric method that assesses differences in central tendency without assuming normality. Two-sided tests were performed for each condition pair using Python’s SciPy package, with significance defined at α = 0.05.

We computed the bootstrapped difference in means between groups by resampling (with replacement) from each distribution independently. Specifically, 10,000 bootstrap iterations were performed per comparison. In each iteration, new samples were drawn from each condition, and the mean difference was recorded. The resulting bootstrap distribution of differences provided an empirical estimate of the effect size and its variability. From this distribution, we computed the 95% confidence interval (CI) using the 2.5th and 97.5th percentiles. A difference was considered statistically significant if the CI did not include zero. This approach allowed robust comparison of condition effects while avoiding parametric assumptions about underlying distributions or equal variances.

### Tissue Processing

To assess viral expression and activity-dependent c-Fos induction, animals were deeply anesthetized and transcardially perfused with phosphate-buffered saline (PBS), followed by 4% paraformaldehyde (PFA). Brains were extracted and post-fixed in 4% PFA for 24 hours at 4°C, then transferred to PBS containing 0.02% sodium azide for storage. Coronal brain sections (50 μm) were cut on a vibratome (Leica VT1000S) in cold PBS using a fresh blade and transferred directly to labeled 12-well plates using a fine paintbrush. Slices were collected in sequential series spanning the anterior-posterior extent of the OFC and stored at 4°C.After 3 washes, slices were mounted onto glass slides with an antifade mounting medium and coverslipped. Images were acquired using an epifluorescence or confocal microscope to confirm location of the GRIN lens.

## Supplementary Figures

**Supplementary Figure 1.**
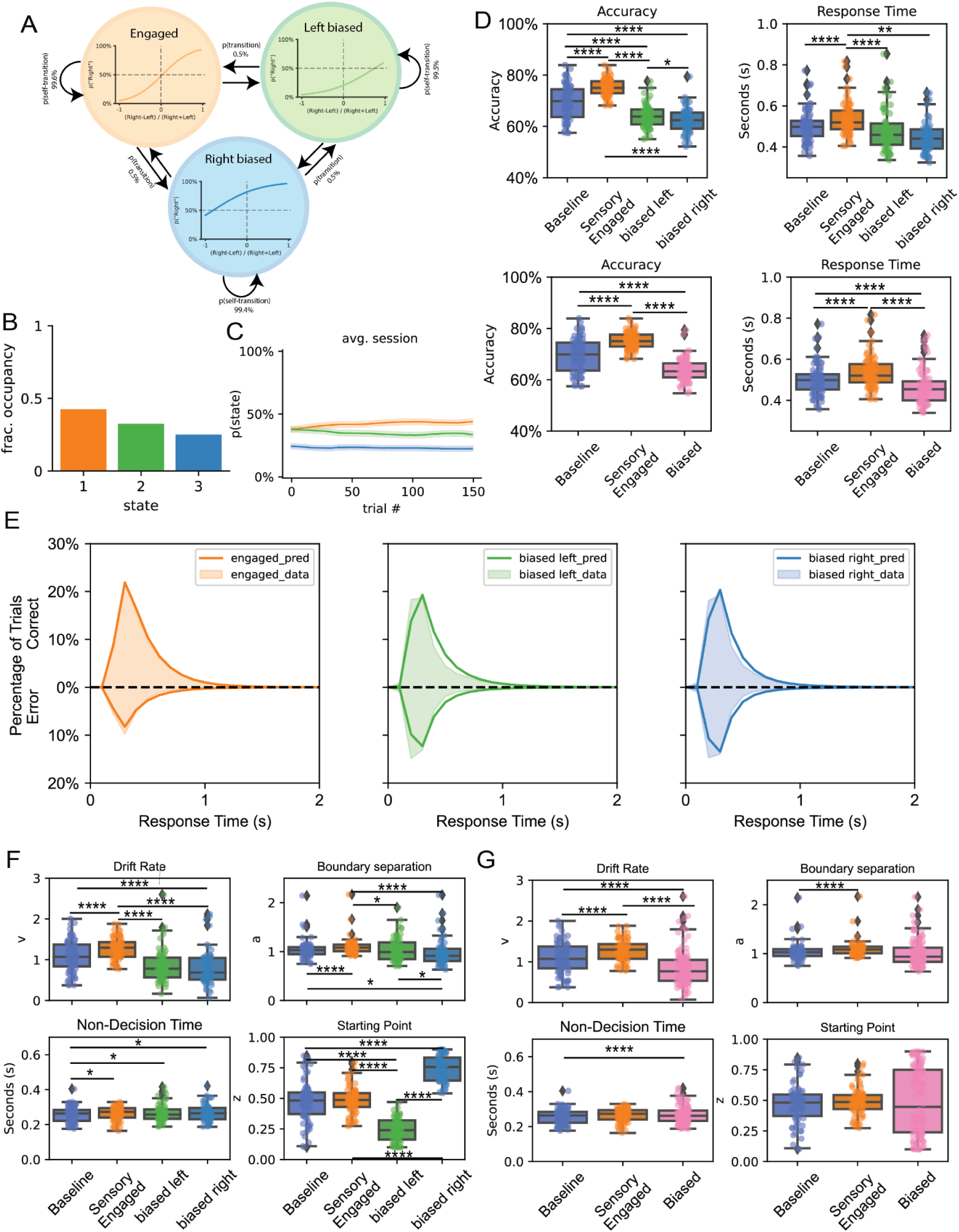
Latent state structure reveals distinct decision policies across biased and engaged strategies. (A) Schematic of the 3-state GLM-HMM model used to infer latent decision-making states: a stimulus-driven (engaged) state and two prior-biased states (left- and right-biased), each associated with distinct strategies. (B) Fraction of time spent in each latent state across all animals, showing dominant occupancy of the engaged state.(C) Probability of occupying each state across trials, averaged across animals and sessions, indicating consistent state dynamics throughout the session. (D) Accuracy (left) and response time (right) for trials classified by latent state identity, showing higher accuracy and longer RTs in the engaged state compared to biased states. Bottom panels split trials combining biased states. (E) Distribution of response times for each latent state, comparing predicted (solid lines) and empirical data (dashed lines), revealing longer and more symmetric RT distributions in the engaged state. (F) Drift diffusion modeling parameters fit to choice-RT data separated by latent state (engaged, left-biased, right-biased): the engaged state is associated with higher boundary separation and reduced starting point bias. (G) Drift diffusion modeling parameters for GLM-HMM states combining biased states. Notations: * = p<0.05, ** = p<0.01, *** = p<0.005, **** = p<0.001.

**Supplementary Figure 2.**
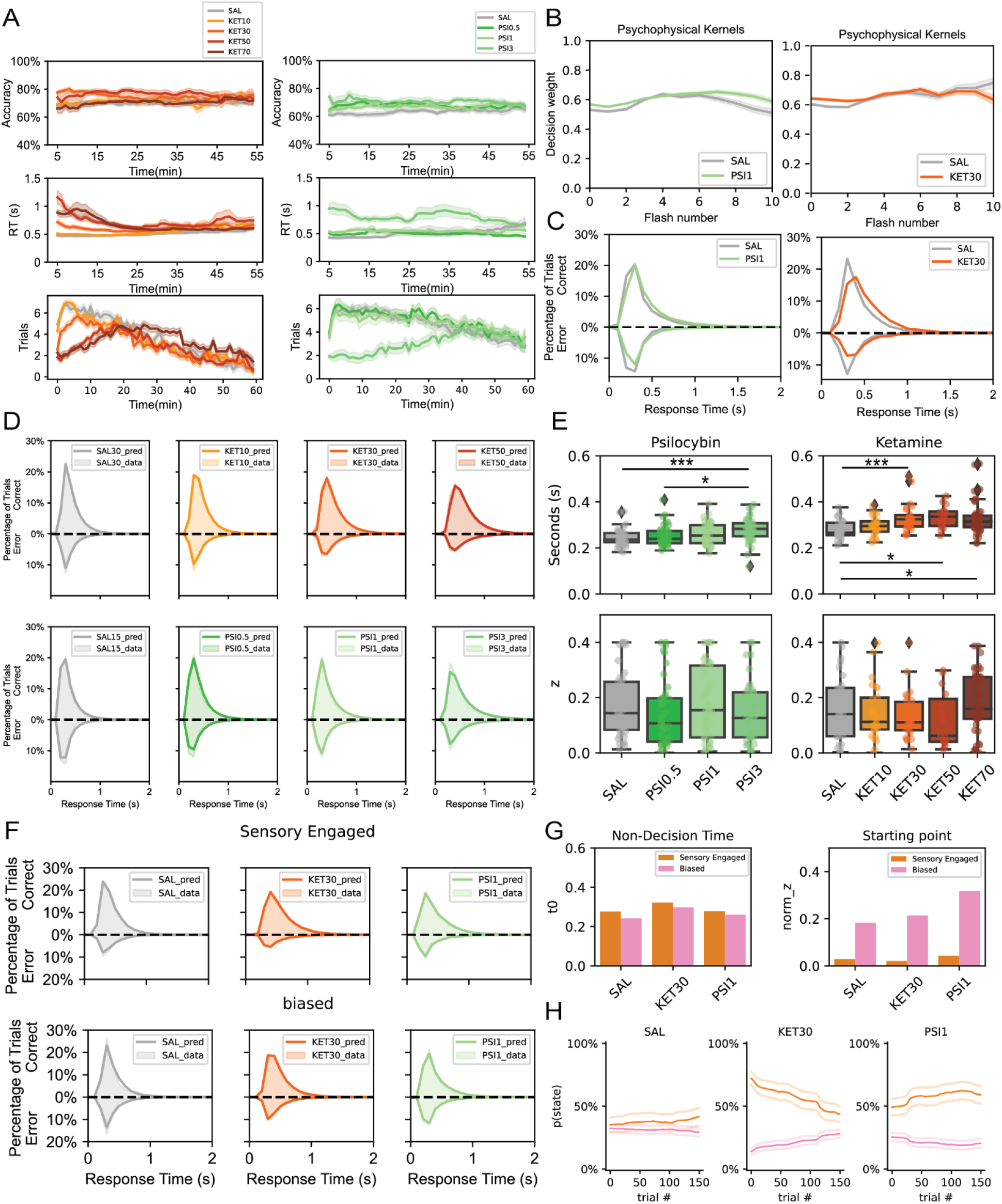
Time-resolved behavior, psychophysical kernels, and latent state modeling support sensory engagement under psychedelics. (A) Rolling averages (5-minute bins) of task performance over a 60-minute session for accuracy (top), response time (middle), and trial rate (bottom) for ketamine (left) and psilocybin (right) conditions, plotted by dose. Shaded areas represent SEM. (B) Psychophysical kernels showing flash weight as a function of flash number across trials under saline, psilocybin (left), and ketamine (right). Flash weights remain consistent across time bins, suggesting stable temporal integration strategies. (C) Histograms of response times for correct and incorrect trials across saline, psilocybin, and ketamine conditions. Psychedelics increase the skew and separation between correct and incorrect RT distributions. (D) DDM model fits to response time distributions across doses for each drug. Solid lines represent empirical data, dashed lines represent DDM fits. Models capture increased RT skew under higher doses of ketamine and psilocybin. (E) DDM parameter estimates for non-decision time (left) and starting point bias (right) across psilocybin and ketamine treatment groups. Ketamine and psilocybin at the highest dose increase non-decision time. (F) DDM predictions for error and correct trials stratified by GLM-HMM state (Sensory-Engaged vs. Biased). (G) Average non-decision time and starting point parameter estimates by GLM-HMM state and treatment. Biased states show higher starting point bias and longer non-decision time. (H) GLM-HMM posterior state probabilities across trials under saline, ketamine 30 mg/kg, and psilocybin 1 mg/kg conditions. Psychedelics increase the occupancy of sensory-engaged states throughout the session.

**Supplementary Figure 3.**
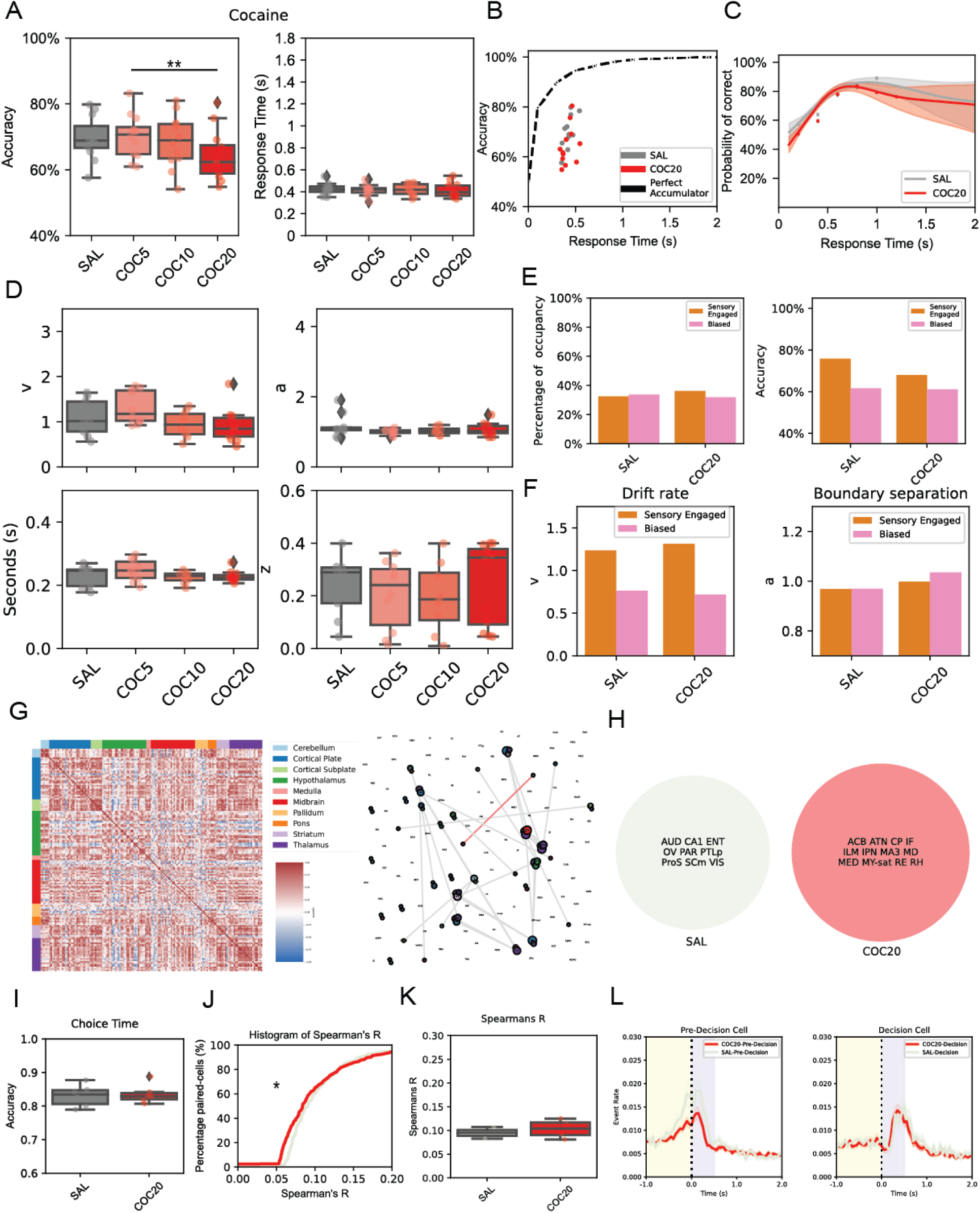
Cocaine does not improve perceptual decision-making or enhance sensory engagement. (A) Behavioral accuracy and response times following acute cocaine administration at varying doses (5, 10, 20 mg/kg). Cocaine does not affect accuracy or response times. (B) Speed-accuracy tradeoff curves comparing saline and cocaine 20 mg/kg to a simulated perfect accumulator. (C) Accuracy as a function of response time for saline and cocaine 20 mg/kg, showing overlapping curves and limited impact of cocaine on evidence integration over time. (D) Drift diffusion model (DDM) parameter estimates for saline and each cocaine dose. Cocaine did not significantly alter drift rate (v), boundary separation (a), non-decision time (t), or starting point (z). (E) State occupancy and accuracy based on GLM-HMM latent state modeling. Cocaine failed to increase occupancy of the sensory-engaged state or improve performance in either state. (F) DDM fit parameters for sensory-engaged and biased states under saline and cocaine 20 mg/kg conditions, showing minimal differences in drift rate and boundary separation. (G) Left: Whole-brain correlation matrix of c-Fos activity across brain regions for cocaine 20 mg/kg during the task. Right: Network graphs illustrate cocaine-induced shifts in network topology and inter-regional connectivity, highlighting altered hubs in the cocaine 20 mg/kg condition. (H) Circle plot showing brain regions with c-fos density significantly different in saline (left, grey) versus cocaine 20 mg/kg (right, red) conditions. (I) Logistic regression decoding accuracy for predicting choice time from OFC neural activity. Cocaine does not significantly enhance decision timing predictability. (J) Histogram of Spearman correlation coefficients among decision-selective OFC neurons. (K) Per-animal averages of Spearman’s R. Cocaine does not reduce inter-neuronal temporal correlation. (L) Trial-aligned neural activity of pre-decision and decision cell populations under saline and cocaine 20 mg/kg conditions (Both: n = 513 neurons, Pre-Decision: n = 314 neurons, Decision: n = 199 neurons). Cocaine does not delay or shift response dynamics in the OFC.

**Supplementary Figure 4:**
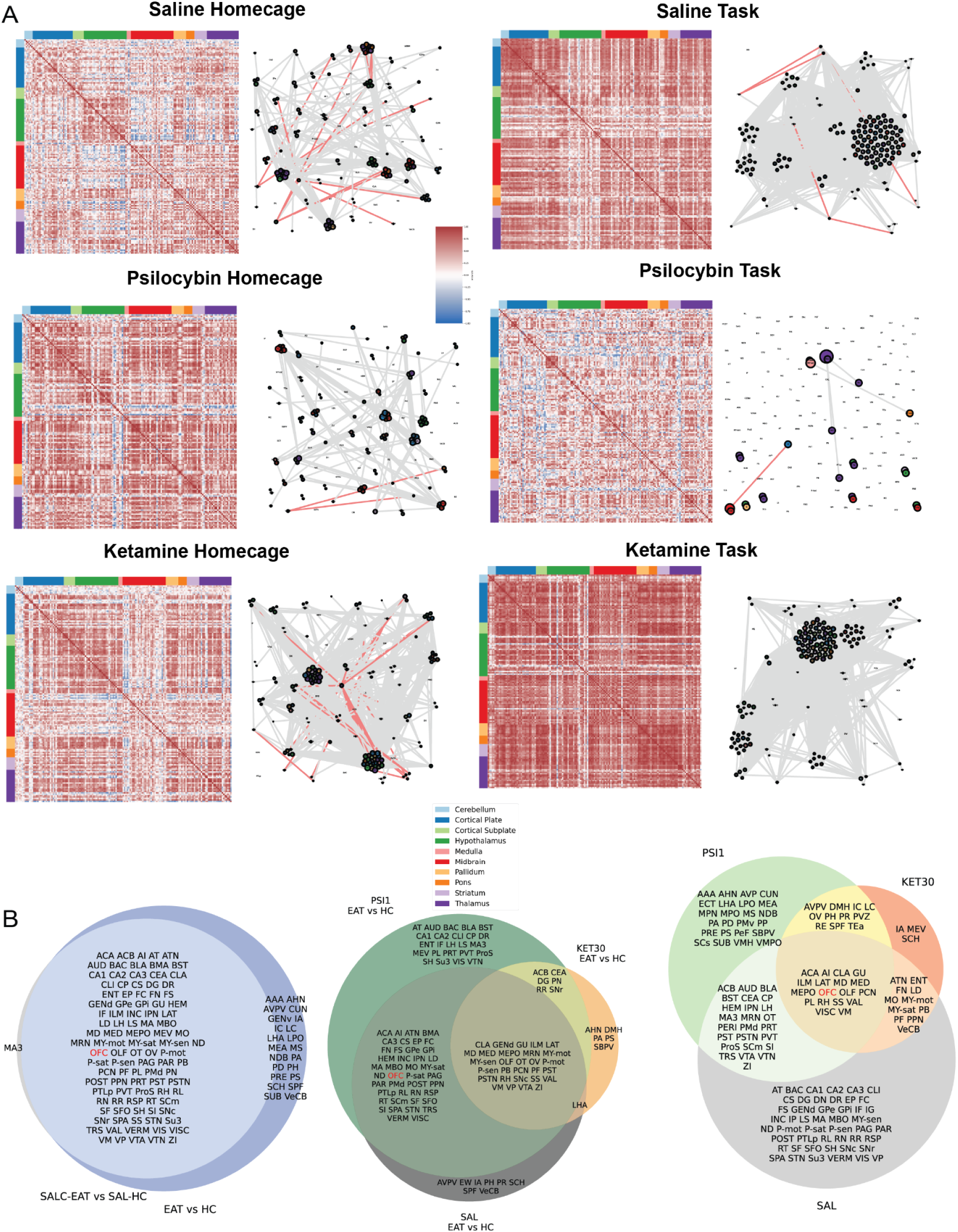
Whole-brain network reorganization under psychedelics during task and homecage conditions.(A) Pairwise Pearson correlation matrices (left) and derived network graphs (right) for c-Fos expression across brain regions under saline, psilocybin, and ketamine, in both homecage (left column) and task (right column) conditions. Each matrix displays region-by-region correlations in c-Fos density (color-coded by anatomical domain), with red representing strong positive correlation and blue representing negative correlation. Network graphs are thresholded based on high-magnitude correlations; edge thickness corresponds to strength, and red lines denote negative correlations. Psychedelics induce large-scale restructuring of network connectivity, with notable reductions in inter-regional coherence during task performance. (B) Venn diagrams illustrate the overlap of significantly modulated regions (based on significant c-fos density differences) across task (EAT) vs. homecage (HC) comparisons. Left: overlap in saline groups for task vs. homecage conditions. Center: overlap in differentially activated regions between psilocybin and ketamine during task vs. homecage comparisons, relative to saline. Right: compound-specific modulations in task-related reorganization (EAT vs. SAL) under psilocybin and ketamine. Psychedelics modulate partially overlapping but distinct subnetworks across forebrain, midbrain, and hindbrain regions during active behavior.

**Supplementary Figure 5:**
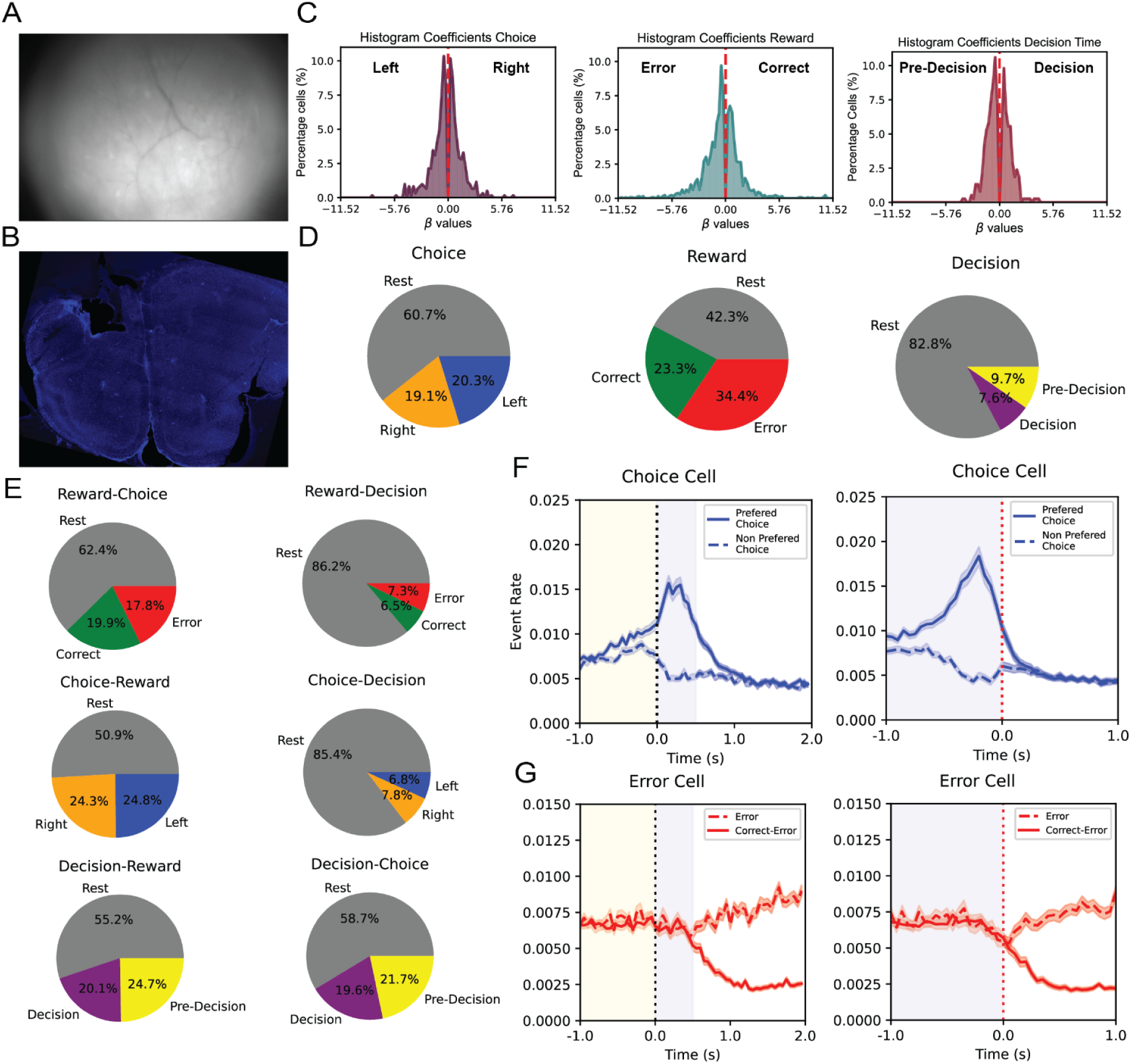
OFC neurons encode decision variables related to choice, reward and decision timing during perceptual evidence accumulation. (A) Representative calcium imaging field of view from the OFC. (B) Histological confirmation of viral expression and implant location in the OFC. (C) Distribution of beta weights (regression coefficients) across neurons for each variable, indicating distinct populations selective for left/right choice, error/correct reward, and pre-decision/decision timing. (D) Proportion of neurons significantly encoding each variable. Choice-selective neurons made up ∼40% of the population (n = 1087 neurons; split between left (n = 560 neurons) and right (n = 527 neurons)), reward-selective neurons ∼60% (favoring correct (n = 644 neurons) or error (n = 951 neurons) outcomes), and ∼18% were selective for decision timing (split between pre-decision ramping (n = 267 neurons) and decision-aligned (n = 209 neurons) activity). (E) Pie charts illustrating the proportion of OFC neurons classified by pairwise overlaps in regression-defined selectivity across task variables during the baseline condition. Each chart represents how many neurons identified as selective for one variable (e.g., Choice) were also selective for another (e.g., Reward), and which specific type (e.g., Left, Right, Error, Correct, etc.). For instance, the top left pie shows how many reward-selective neurons were also selective for choice direction, and whether they encoded left, right, or rest epochs. These pairwise overlaps reveal how mixed selectivity and task-stage encoding are distributed across the OFC population. (F-G) Average calcium event rate of choice-selective (F) and reward-selective (G) neurons aligned to trial initiation(left) and choice (right). Choice cells activated for preferred options and suppressed for non-preferred ones prior to decision commitment. Reward cells encoded outcome selectivity post-choice.

**Supplementary Figure 6.**
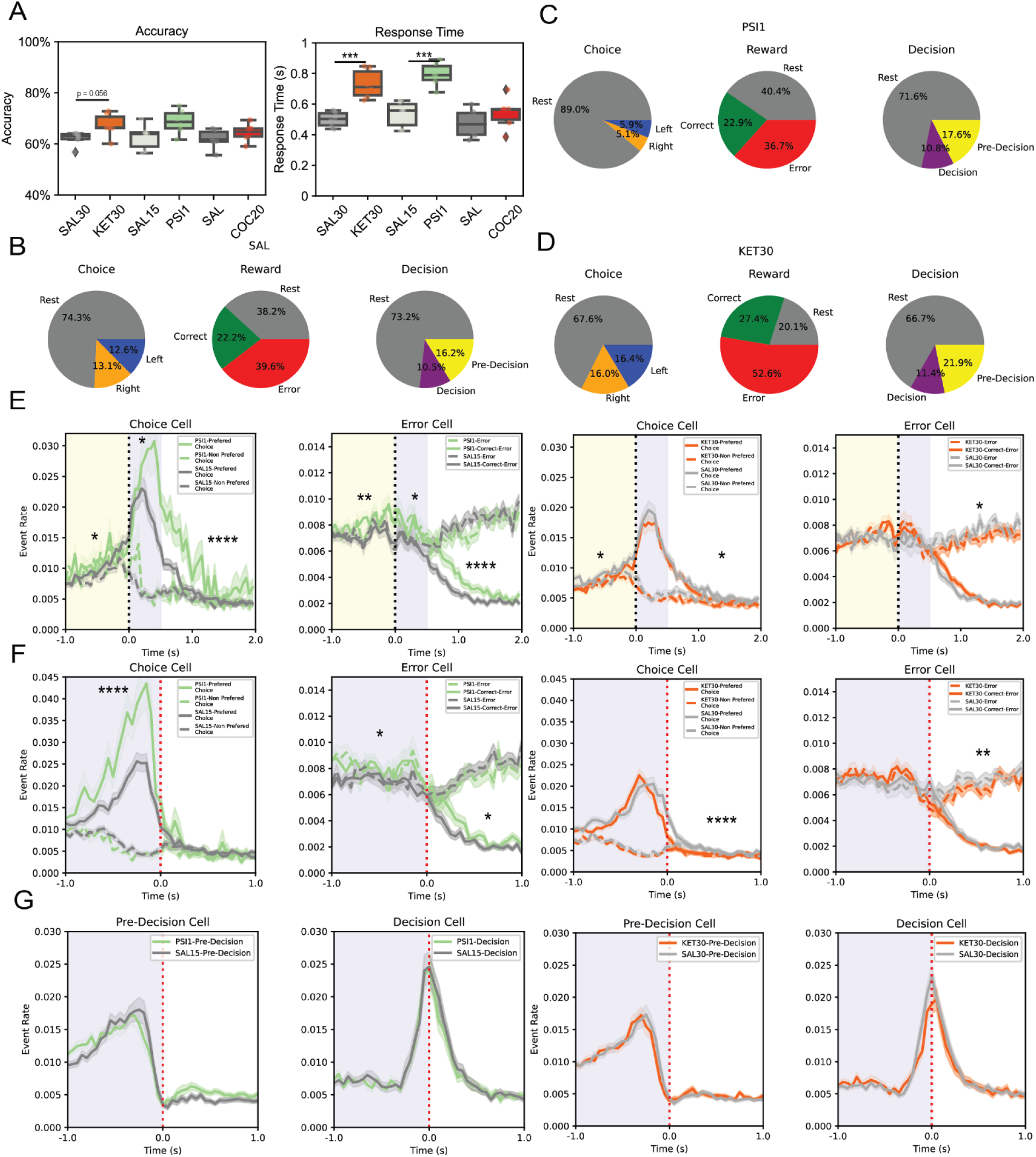
Cell-type classifications and temporal dynamics of OFC neurons across drug conditions. (A) Behavioral performance across experimental groups in Thy1-GCaMP6f mice, showing average accuracy (left) and response time (right) for saline, ketamine, psilocybin, and cocaine treatment conditions. (B-D) Pie charts of OFC cell classifications under saline (B), psilocybin 1 mg/kg (C), and ketamine 30 mg/kg (D), showing distributions of Choice, Reward, and Decision-selective neurons. (E) Peri-event time histograms (PETHs) for choice- and error-selective cells under psilocybin 1 mg/kg and ketamine 30 mg/kg compared to their saline controls, aligned to trial initiation (dotted black line). Shading represents SEM; vertical colored boxes highlight pre-initiation and trial windows. Psilocybin enhanced early and sustained choice-selective activity; ketamine prolonged error-related responses. (F) PETHs comparing activity of choice- and error-related cells between compound and saline conditions, aligned to the choice moment (red dashed line). Psilocybin increased choice signal encoding while dampening error cell responses. Ketamine increased late error cells during error trials activity. (G) PETHs of pre-decision and decision-selective cells, showing response patterns relative to choice timing (red dashed line).

## References

1. Carhart-Harris, R. L. et al. Psilocybin with psychological support for treatment-resistant depression: six-month follow-up. Psychopharmacology (Berl.) 235, 399–408 (2018).

2. Feder, A. et al. A Randomized Controlled Trial of Repeated Ketamine Administration for Chronic Posttraumatic Stress Disorder. Am. J. Psychiatry 178, 193–202 (2021).

3. Carhart-Harris, R. L. & Friston, K. J. REBUS and the Anarchic Brain: Toward a Unified Model of the Brain Action of Psychedelics. Pharmacol. Rev. 71, 316–344 (2019).

4. Vollenweider, F. X. & Preller, K. H. Psychedelic drugs: neurobiology and potential for treatment of psychiatric disorders. Nat. Rev. Neurosci. 21, 611–624 (2020).

5. Brunton, B. W., Botvinick, M. M. & Brody, C. D. Rats and humans can optimally accumulate evidence for decision-making. Science 340, 95–98 (2013).

6. Scott, B. B. et al. Fronto-parietal Cortical Circuits Encode Accumulated Evidence with a Diversity of Timescales. Neuron 95, 385–398.e5 (2017).

7. Scott, B. B., Constantinople, C. M., Erlich, J. C., Tank, D. W. & Brody, C. D. Sources of noise during accumulation of evidence in unrestrained and voluntarily head-restrained rats. eLife 4, e11308 (2015).

8. Gupta, D., DePasquale, B., Kopec, C. D. & Brody, C. D. Trial-history biases in evidence accumulation can give rise to apparent lapses in decision-making. Nat. Commun. 15, 662 (2024).

9. Chakravarty, S. et al. A cross-species framework for investigating perceptual evidence accumulation. eLife 14, (2025).

10. Kane, G. A., Senne, R. A. & Scott, B. B. Rat movements reflect internal decision dynamics in an evidence accumulation task. J. Neurophysiol. 132, 1608–1620 (2024).

11. Schuck, N. W., Cai, M. B., Wilson, R. C. & Niv, Y. Human Orbitofrontal Cortex Represents a Cognitive Map of State Space. Neuron 91, 1402–1412 (2016).

12. Wilson, R. C., Takahashi, Y. K., Schoenbaum, G. & Niv, Y. Orbitofrontal Cortex as a Cognitive Map of Task Space. Neuron 81, 267–279 (2014).

13. Ashwood, Z. C. et al. Mice alternate between discrete strategies during perceptual decision-making. Nat. Neurosci. 25, 201–212 (2022).

14. Ratcliff, R. & McKoon, G. The Diffusion Decision Model: Theory and Data for Two-Choice Decision Tasks. Neural Comput. 20, 873–922 (2008).

15. Bogacz, R., Wagenmakers, E.-J., Forstmann, B. U. & Nieuwenhuis, S. The neural basis of the speed-accuracy tradeoff. Trends Neurosci. 33, 10–16 (2010).

16. Frank, M. J. Hold your horses: a dynamic computational role for the subthalamic nucleus in decision making. Neural Netw. Off. J. Int. Neural Netw. Soc. 19, 1120–1136 (2006).

17. Cavanagh, S. E., Lam, N. H., Murray, J. D., Hunt, L. T. & Kennerley, S. W. A circuit mechanism for decision-making biases and NMDA receptor hypofunction. eLife 9, e53664 (2020).

18. Park, A. J. et al. Reset of hippocampal–prefrontal circuitry facilitates learning. Nature 1–5 (2021) doi:10.1038/s41586-021-03272-1.

19. Renier, N. et al. Mapping of Brain Activity by Automated Volume Analysis of Immediate Early Genes. Cell 165, 1789–1802 (2016).

20. Hunt, L. T. et al. Mechanisms underlying cortical activity during value-guided choice. Nat. Neurosci. 15, 470–476 (2012).

21. Lak, A. et al. Dopaminergic and Prefrontal Basis of Learning from Sensory Confidence and Reward Value. Neuron 105, 700–711.e6 (2020).

22. Kepecs, A., Uchida, N., Zariwala, H. A. & Mainen, Z. F. Neural correlates, computation and behavioural impact of decision confidence. Nature 455, 227–231 (2008).

23. Tibshirani, R. Regression Shrinkage and Selection Via the Lasso. J. R. Stat. Soc. Ser. B Stat. Methodol. 58, 267–288 (1996).

24. Ecker, A. S. et al. Decorrelated Neuronal Firing in Cortical Microcircuits. Science 327, 584–587 (2010).

25. Averbeck, B. B., Latham, P. E. & Pouget, A. Neural correlations, population coding and computation. Nat. Rev. Neurosci. 7, 358–366 (2006).

26. Deco, G. et al. Whole-Brain Multimodal Neuroimaging Model Using Serotonin Receptor Maps Explains Non-linear Functional Effects of LSD. Curr. Biol. CB 28, 3065–3074.e6 (2018).

27. Luppi, A. I. et al. LSD alters dynamic integration and segregation in the human brain. NeuroImage 227, 117653 (2021).

28. Timmermann, C. et al. Neural correlates of the DMT experience assessed with multivariate EEG. Sci. Rep. 9, 16324 (2019).

29. Preller, K. H. et al. Effective connectivity changes in LSD-induced altered states of consciousness in humans. Proc. Natl. Acad. Sci. U. S. A. 116, 2743–2748 (2019).

30. Mason, N. L., Dolder, P. C. & Kuypers, K. P. Reported effects of psychedelic use on those with low well-being given various emotional states and social contexts. Drug Sci. Policy Law 6, 2050324519900068 (2020).

31. Yu, Z. et al. Alterations in brain network connectivity and subjective experience induced by psychedelics: a scoping review. Front. Psychiatry 15, (2024).

32. Preller, K. H. et al. Psilocybin Induces Time-Dependent Changes in Global Functional Connectivity. Biol. Psychiatry 88, 197–207 (2020).

33. Sul, J. H., Kim, H., Huh, N., Lee, D. & Jung, M. W. Distinct roles of rodent orbitofrontal and medial prefrontal cortex in decision making. Neuron 66, 449–460 (2010).

34. Moreno-Bote, R. et al. Information-limiting correlations. Nat. Neurosci. 17, 1410–1417 (2014).

35. Kanitscheider, I., Coen-Cagli, R. & Pouget, A. Origin of information-limiting noise correlations. Proc. Natl. Acad. Sci. 112, E6973–E6982 (2015).

36. Carhart-Harris, R. L. et al. Neural correlates of the LSD experience revealed by multimodal neuroimaging. Proc. Natl. Acad. Sci. 113, 4853–4858 (2016).

37. Davoudian, P. A., Shao, L.-X. & Kwan, A. C. Shared and Distinct Brain Regions Targeted for Immediate Early Gene Expression by Ketamine and Psilocybin. http://biorxiv.org/lookup/doi/10.1101/2022.03.18.484437 (2022) doi:10.1101/2022.03.18.484437.

38. Aboharb, F. et al. Classification of psychedelics and psychoactive drugs based on brain-wide imaging of cellular c-Fos expression. Nat. Commun. 16, 1590 (2025).

39. Dhawale, A. K. et al. Automated long-term recording and analysis of neural activity in behaving animals. eLife 6, e27702 (2017).

40. Poddar, R., Kawai, R. & Ölveczky, B. P. A Fully Automated High-Throughput Training System for Rodents. PLOS ONE 8, e83171 (2013).

41. Wood, S. N., Li, Z., Shaddick, G. & Augustin, N. H. Generalized Additive Models for Gigadata: Modeling the U.K. Black Smoke Network Daily Data. J. Am. Stat. Assoc. 112, 1199–1210 (2017).

42. Vallat, R. Pingouin: statistics in Python. J. Open Source Softw. 3, 1026 (2018).

43. Churchland, A. K., Kiani, R. & Shadlen, M. N. Decision-making with multiple alternatives. Nat. Neurosci. 11, 693–702 (2008).

44. Nelder, J. A. & Wedderburn, R. W. M. Generalized Linear Models. J. R. Stat. Soc. Ser. Gen. 135, 370–384 (1972).

45. Linderman, S. et al. Bayesian Learning and Inference in Recurrent Switching Linear Dynamical Systems. in *Proceedings of the 20th International Conference on Artificial Intelligence and Statistics* 914–922 (PMLR, 2017).

46. Ratcliff, R. A theory of memory retrieval. Psychol. Rev. 85, 59–108 (1978).

47. Park, Y.-G. et al. Protection of tissue physicochemical properties using polyfunctional crosslinkers. Nat. Biotechnol. 37, 73–83 (2019).

48. Vousden, D. A. et al. Whole-brain mapping of behaviourally induced neural activation in mice. Brain Struct. Funct. 220, 2043–2057 (2015).

49. Dorst, K. E. et al. Hippocampal Engrams Generate Variable Behavioral Responses and Brain-Wide Network States. J. Neurosci. 44, (2024).

